## Supporting Information for "Determining accurate conformational ensembles of intrinsically disordered proteins at atomic resolution"

Kaushik Borthakur,[1] Thomas R. Sisk,[1] Francesco P. Panei ,[2] Massimiliano Bonomi,[2] and Paul Robustelli*,[1]

[1] *Dartmouth College, Department of Chemistry, Hanover, NH, 03755, USA*

[2] *Institut Pasteur, Université Paris Cité, CNRS UMR 3528, Computational Structural Biology Unit, Paris, France*

*

### Theory

In the following sections we provide a brief overview of the theory of the maximum entropy reweighting and the treatment of different sources error with regularization parameters. We focus on the formalism proposed by Bussi et al.[1–3] which is the basis of our approach.

#### Overview of maximum entropy reweighting

A MD simulation generates a prior distribution $P_0(x)$ of conformations $x$. In initial implementations,[2,4] maximum entropy reweighting approaches were designed to determine the distribution $P_1(x)$ which is the closest to $P_0(x)$ that fits *exactly* each experimental data point $f_i^{exp}$. To measure the statistical distance between the two distributions $P_0(x)$ and $P_1(x)$, one can use the Kullback–Leibler divergence, defined as:

$$D_{KL}[P_1|P_0] = \int P_1(x) \ln \frac{P_1(x)}{P_0(x)} dx \tag{1}$$

To determine $P_1(x)$, the closest distribution to $P_0(x)$ that fits the ensemble-averaged data, the KL divergence is then minimized subject to the following two constraints:

$$\int P_1(x) f_i(x) dx - f_i^{exp} = 0 \text{ for } i = 0, 1, 2, ...., M \tag{2}$$

$$\int P_1(x) dx = 1 \tag{3}$$

where $f_i(x)$ is the forward model needed to predict the value of observable $i$ for each conformation $x$. In Eq. 2, each experimental observation constrains the ensemble average of the corresponding forward model $f_i(x)$ computed over the distribution $P_1(x)$ to be exactly equal to $f_i^{exp}$. The second constraint in Eq. 3 ensures that $P_1(x)$ is normalized.

The problem of finding the distribution $P_1(x)$ that is the best representation of the state of knowledge of a system after observing a set of experimental data can be cast in the framework of information theory. In this framework, amongst all the possible distributions compatible with the data, the one that has the highest degree of uncertainty should be selected in order to ensure minimal bias and that the data has been used as conservatively as possible. This uncertainty is expressed by the Shannon entropy[5] of the random variable $x$ with distribution $P_1(x)$ with respect to the reference distribution $P_0(x)$:

$$S[P_1|P_0] = -\int P_1(x) \ln \frac{P_1(x)}{P_0(x)} dx = -D_{KL}[P_1|P_0] \tag{4}$$

Maximizing the Shannon entropy in Eq. 4 to determine $P_1(x)$ is thus equivalent to minimizing the KL divergence in Eq. 1, hence the name of maximum entropy reweighting.

The solution for the minimization of the KL divergence subject to the constraints in Eq. 2 and 3 can be achieved by using Lagrange multipliers. We first define the following Lagrange function:

$$\mathcal{L} = D_{KL}[P_1|P_0] - \sum_{i=1}^{M} \lambda_i \left[ \int P_1(x) f_i(x) dx - f_i^{exp} \right] - \mu \left[ \int P_1(x) dx - 1 \right] \tag{5}$$

where $\lambda_i$ and $\mu$ are Lagrange multipliers. There is one Lagrange multiplier $\lambda_i$ for each of the of the $M$ experimental data points being used as constraints. To find stationary points of the Lagrangian, we set $\partial \mathcal{L} / \partial P_1 = 0$ and neglect the normalization factor. This leads to the following form for the desired distribution $P_1(x)$:

$$P_1(x) \propto e^{-\sum_{i=1}^{M} \lambda_i f_i(x)} P_0(x) \tag{6}$$

Approaches such as gradient descent or appropriate quasi-Newton methods can be utilized to calculate the values of the Lagrange multipliers, which ultimately yield to a new set of weights, one for each conformation of the ensemble:

$$w(x) \propto e^{-\sum_{i=1}^{M} \lambda_i f_i(x)} \tag{7}$$

#### Introducing different source of errors

By fitting exactly the experimental observations, the formalism introduced above neglects the fact that experimental data as well as forward models can be affected by random and systematic errors and therefore data should not be matched *exactly* to avoid overfitting to noise. In order to alleviate the problem of data overfitting, maximum entropy reweighting approaches can be modified to account for different sources of errors as well as a limited confidence in the prior distribution. Different approaches have been proposed over the years.[2,6–9] In the formalism of Bussi et al.,[2] errors are modeled by modifying the constraints of Eq. 2 by introducing an auxiliary variable for each data point $\epsilon_i$, which represents the difference between experimental and predicted values. The new constraints are hence defined as follows:

$$\int P_1(x)\,(f_i(x) + \epsilon_i)\,dx - f_i^{exp} = 0 \text{ for } i = 0, 1, 2, ...., M \tag{8}$$

Errors can be modeled by choosing a proper prior distribution function for the variables $\epsilon_i$. We choose a Gaussian prior with fixed standard deviation $\sigma_i$ for the $i^{th}$ observable.

$$P_0(\epsilon_i) \propto \prod_{i=1}^{M} e^{-\left(\frac{\epsilon_i^2}{2\sigma_i^2}\right)} \tag{9}$$

The value of $\sigma_i$ corresponds to the level of confidence in the $i^{th}$ experimental data point and ultimately determines how well the reweighted ensemble will fit the data. $\sigma_i = \infty$ implies no confidence in the data and results in a reweighted ensemble that is identical to the unbiased simulation. $\sigma_i = 0$ implies complete confidence in the data and results in a reweighted ensembles in which the $i^{th}$ data point is exactly matched, as in Eq. 2. The selection of appropriate values of the regularization parameters $\sigma_i$ for each experimental data point is therefore essential to balance one's confidence in a prior model (ie. the accuracy of a force field), error estimates of experimental measurements and error estimates of the accuracy of forward models for predicting experimental data. Ultimately, the properties of the resulting conformational ensembles will be highly dependent on the relative magnitudes of the regularization parameters selected for each experimental data point.

Different strategies have been developed to identify the optimal values of $\sigma_i$. Typically, when performing reweighting using multiple types of experimental data, one value of $\sigma_i$ is assigned to each class of experimental data, such as SAXS intensities or NMR chemical shifts, RDCs or scalar couplings. A grid search is then performed, usually independently for each class of experimental data, and the optimal value of $\sigma_i$ is identified for each class to maximize the agreement of the reweighted ensemble with independent experimental data not used for reweighting.[10] A final reweighting step using all the available experimental data is then performed.

#### Determining regularization parameters

In the approach proposed in this work, we assign one regularization parameter $\sigma_i$ to each data point. To optimize the grid search over the space of $\sigma_i$, using a similar functional form to previous work[11] we decompose this parameter into two contributions:

$$\sigma_i = \sqrt{\sigma_{i,MD}^2 + \sigma_{reg}^2}. \tag{10}$$

where $\sigma_{reg}$ is the regularization parameter, one per data type, that describes both experimental and forward model errors of this specific type of experiment. The $\sigma_{i,MD}$ parameter, one per data point, represents both the statistical errors of forward models to predict experimental data and statistical errors that result from calculating average quantities over a finite-size ensemble. We calculate $\sigma_{i,MD}$ using the Flyvbjerg block analysis[11,12] technique to account for the correlation between adjacent frames in the MD simulation or the standard error of the mean for uncorrelated data. We use a single free parameter, the (normalized) Kish effective sample size,[3,13] or *Kish ratio* $K$, to determine the optimal values of $\sigma_{reg}$ for each experimental data type as described in the main text. $K$ is defined as

$$K = \frac{1}{N}\left(\sum_{i=1}^{N} w_i\right)^2 / \sum_{i=1}^{N} w_i^2 \tag{11}$$

where $w_i$ is value of the statistical weight of the $i$th frame.

### Developing a maximum entropy reweighting protocol with a single free parameter

Here, we summarize the key features of our approach and provide illustrative examples that highlight the rationale behind the choices made. We build upon the formalism of Bussi and coworkers,[1–3] where the strength of the restraint on each experimental data point $i$ is determined by the value of an adjustable regularization parameter $\sigma_i$ (Eq. 10). A key feature of our reweighting procedure is the use of effective ensemble sizes of reweighted ensembles to select the values of regularization parameters for reweighting. We quantify the effective ensemble size of reweighted ensemble using the Kish ratio[3,13] (Eq. 11). Briefly, the Kish ratio ($K$) is a measure of the fraction of frames of an original ensemble with significant statistical weights after reweighting. A reweighted ensemble where ∼90% of the frames have statistical weights close to zero will have a Kish ratio of $K \approx 0.10$. Conversely, if an unbiased MD ensemble contains 10,000 frames, a reweighted ensemble with a Kish ratio $K \approx 0.01$ would contain approximately 100 frames with statistical weights substantially larger than zero.

We initially attempted to reweight IDP ensembles using a cross-validation approach to determine the relative strengths of restraints for different experimental data types. We performed reweighting using a single value of the regularization parameter $\sigma$ for all experimental data points of a given type (ie. C$\alpha$ chemical shifts, C$\beta$ chemical shifts, SAXS intensities) and iteratively decreased the value of $\sigma$ while monitoring: i) the root mean squared error (RMSE) between calculated and experimental data used as restraints; ii) the RMSE between calculated end experimental cross-validating data not used as restraints; iii) the Kish ratio of the reweighted ensemble. We sought to identify an optimal value of $\sigma$ for each experimental data type corresponding to the point where further reducing $\sigma$ caused agreement with unrestrained cross-validating experimental data to deteriorate. We found, however, that many experimental data types were highly correlated and the RMSE of several cross-validating data sets continued to improve as the Kish ratio approached zero (SI Figure 1).

We tested several Bayesian and heuristic approaches to infer the optimal values of regularization parameters for each type of experimental data by monitoring the agreement of restrained and unrestrained cross-validation data using empirical RMSE thresholds and L-curve or "elbow method" analyses.[6,10,14] In many instances, we observed that these approaches produced reweighted ensembles with extremely small effective ensemble sizes, where only hundreds of structures from an initial MD ensemble of nearly 30,000 structures maintained appreciable statistical weights after reweighting ($K < 0.01$). When reweighting ensembles derived from different force fields, we observed that these approaches produced reweighted ensembles with dramatic differences in their effective ensemble size, depending on the initial agreement of unbiased MD ensembles with experimental data. This complicated global comparisons of the accuracy and structural properties of reweighted ensembles derived from different force fields.

As we desire reweighted ensemble sizes that are large enough to provide reasonable statistics for the conformational properties of IDPs, such as the overall topology of backbone conformations and the relative positions of sidechains, we opted to utilize the value of the Kish ratio of reweighted ensembles to determine regularization parameters for each data type. To account for the errors of forward models used to calculate experimental data and the effects of calculating average quantities over a finite-size ensemble, we introduced an additional regularization parameter $\sigma_{i,MD}$ for each experimental data point $i$ used for reweighting (Eq. 10). Inspired by previous work,[11] We calculated $\sigma_{i,MD}$ by applying the Flyvbjerg block analysis method[12] to the forward model predictions obtained from the unbiased MD trajectory.

With this addition, we obtain one regularization parameter for each experimental data point $\sigma_i = \sqrt{\sigma_{i,MD}^2 + \sigma_{reg}^2}$, where $\sigma_{reg}$ is the same for all data of a given data type (ie. $\sigma_{reg-C\alpha}$, $\sigma_{reg-C\beta}$, $\sigma_{reg-SAXS}$).

To determine an optimal minimum Kish ratio threshold for reweighting IDP ensembles, we examined how the conformational properties of reweighted ensembles changed as a function of $\sigma_{reg}$ for each data type. We sought to identify a minimum Kish ratio threshold where the conformational properties of reweighted ensembles, such as secondary structure content, radius of gyration ($R_g$) and populations of intramolecular contacts, remained relatively consistent as $\sigma_{reg}$ was decreased and further improvements in the agreement between calculated and experimental data were largely achieved by increasing the sparsity of the reweighted ensemble. For each data type, as the strength of experimental restraints was increased (by decreasing $\sigma_{reg}$) we generally observed an initial regime where agreement with experimental data was achieved by changing the statistical weights of the structures belonging to the most populated regions of conformational space sampled in unbiased MD simulations. As Kish ratios approached smaller values (ie. $K < 0.10$), improved agreement with experimental data was often achieved by favoring a smaller number of structures from regions of conformational space with unfavorable free energies in unbiased MD simulations.

An illustrative example of this behavior is shown in SI Figure 2. Here, we use only two experimental backbone NMR chemical shifts of a single residue as restraints to reweight the unbiased a99SB-*disp* MD ensemble of PaaA2. We examine how reweighted distributions of the predicted values of these chemical shifts change as we increase the strength of these two restraints. We observe that as we decrease $\sigma_{reg}$ from 16.0 to 2.0, we improve agreement with experimental data by increasing the statistical weights of conformations near the most populated free energy basins of the original unbiased MD ensemble. When we further increase the strength of experimental restraints by setting $\sigma_{reg} = 1.0$, the Kish ratio drops to $K = 0.003$, corresponding to an effective reweighted ensemble size of approximately one hundred structures from an initial ensemble of 29,976 structures. Notably, we observe that very few of the conformations that maintain non-negligible statistical weights in the reweighted ensemble are located within the free energy basins of the original unbiased MD ensemble (SI Figure 2). This dramatic reduction in the effective ensemble size and reshaping of the free energy surface results in an extremely marginal improvement in the agreement between the calculated and experimental values of these chemical shifts relative to the reweighted ensemble obtained with $\sigma_{reg} = 2.0$, which has a Kish ratio $K = 0.11$. This marginal improvement is likely within the noise of chemical shift predictor SPARTA+.[15]

We performed similar analyses for each experimental data type, monitoring how the free energy surfaces of reweighted ensembles and agreement with unrestrained cross-validating data varied with $\sigma_{reg}$. Based on these analyses, we elected to select regularization parameters using a Kish ratio threshold of $K = 0.10$. This means we iteratively decrease $\sigma_{reg}$ for each individual data type until the resulting reweighted ensemble has a Kish ratio $K < 0.10$, and select the smallest value of $\sigma_{reg}$ where $K \geq 0.10$ as the final $\sigma_{reg}$ value for that data type. After establishing an initial $\sigma_{reg}$ for each individual data type, we perform a global reweighting using all experimental data as restraints simultaneously. In order to ensure a suitable ensemble size for the final reweighted ensemble, we rescaled the regularization parameter of each data type by a global scaling factor $\sigma_{reg-Global}$. We iteratively decrease the value of $\sigma_{reg-Global}$ and select the minimum value of $\sigma_{reg-Global}$ that produces a reweighted ensemble with Kish ratio $K \geq 0.10$ as the final value for global reweighting using all experimental data as restraints. The full reweighting protocol is illustrated for the a99SB-*disp* MD ensemble of A$\beta$40 in SI Figure 3. We note that the choice of an optimal Kish ratio threshold is ultimately a subjective decision that will depend on the initial ensemble size of unbiased ensembles, the desired level of agreement with experimental data and the the desired degree of statistical sampling of properties of interest in reweighted ensembles. We found that varying the Kish ratio threshold between $K = 0.05$ and $K = 0.25$ did not appreciably change the structural properties of reweighted IDP ensembles or any of the conclusions of this study.

#### A maximum entropy reweighting protocol with a single free parameter improves agreement with extensive experimental datasets with minimal overfitting

We illustrate the key components of the proposed reweighting procedure by reweighting the unbiased a99SB-*disp* MD ensemble of A$\beta$40 (SI Figure 3). We illustrate the process of determining $\sigma_{reg}$ for each experimental data type of A$\beta$40 in SI Figure 3A and the process of determining the global regularization parameter scaling factor $\sigma_{reg-Global}$ in Figure SI Figure 3B. In both plots, we denote the selected Kish ratio threshold $K = 0.10$ with a dashed line. The proposed automated reweighting procedure simultaneously improves agreement with all types of experimental data in this system without requiring any manual adjustments of the relative strength of experimental restraints. We observe that the RMSE between calculated and experimental data monotonically decreases for all types of experimental data as the effective ensemble size of the reweighted A$\beta$40 ensemble decreases from 30,000 to 3000, corresponding to the selected Kish Ratio threshold of $K$=0.10 (SI Figure 3A). We find that the RMSE between calculated and experimental data continues to monotonically decrease for all types of data until the effective ensemble size reaches ~150 structures ($K$~0.005). This demonstrates that while we select a Kish Ratio threshold of $K$=0.10 to obtain a desired degree of statistical sampling in our final reweighted A$\beta$40 ensemble, we could further improve global agreement with experimental data if we were willing to produce a sparser reweighted ensemble with less statistical sampling of conformational properties of interest. We note the RMSE between calculated and experimental data begins to deteriorate for extremely small effective ensembles sizes containing fewer than 70 structures, which we attribute to instability in the L-BFGS minimizer used to minimize the Lagrange function in SI Eq. 5 in the presence of 237 experimental restraints.

A key feature of the reweighting protocol proposed here is that the relative strengths of restraints for different types of experimental data are determined automatically. It is therefore important to determine if the proposed procedure simultaneously improves agreement with all data types in large experimental NMR and SAXS datasets or if the procedure is overfit to a subset of experimental data. To asses this, we performed i) reweighting using each experimental data type as the only restraint; ii) a global reweighting using all experimental data as restraints; and iii) a leave-one-out cross-validation reweighting using all but one experimental data type as restraints. The results of these reweighting calculations are shown for the a99SB-*disp* MD ensemble of A$\beta$40 in SI Figure 3C and SI Figure 3D. In SI Figure 3D we compare the accuracy of reweighted ensembles to the unbiased ensemble using a normalized RMSE ($\mathrm{RMSE_N}$) where we normalize the value of the RMSE calculated for each experimental data type by the corresponding value in the unbiased a99SB-*disp* MD ensemble ($\mathrm{RMSE_N} = \mathrm{RMSE}/\mathrm{RMSE}_{\mathrm{a99SB}-disp}$).

We observe that when we use all available experimental NMR data ($C\alpha$, $C\beta$, $H\alpha$, $H$, and $N$ backbone NMR chemical shifts, backbone $^3J_{HNH\alpha}$ coupling constants, and backbone RDCs) as restraints our reweighting protocol simultaneously improves agreement with all experimental data types (SI Figure 3D, "All Restraints"). For each experimental data type, we obtain a smaller RMSE between calculated and experimental data when we perform reweighting using only those data as restraints (SI Figure 3D "Individual Restraints") compared to the RMSE obtained when performing reweighting using all available data as restraints. This demonstrates that satisfying multiple types of data requires trade offs in the agreement with each type of restraint and that our reweighting procedure naturally balances the stregnths of different types of restraints without requiring manual adjustment of regularization parameters or cross-validation thresholds.

We observe that the RMSE between calculated and experimental data for each data type withheld in leave-one-out cross validation reweighting calculations is very similar to the RMSE obtained when reweighting using all experimental data as restraints (SI Figure 3D, "Cross validation"). This suggests that using a Kish ratio threshold of $K \geq 0.10$ to determine regularization parameters for each data type results in minimal overfitting when all experimental data are used as restraints in this system. We observe that the structural properties (secondary structure populations, intramolecular contact populations and $R_g$ distribution) of the A$\beta$40 conformational ensemble obtained by reweighting with all experimental data and the structural properties of conformational ensembles obtained from leave-one-out cross-validation reweighting calculations were largely indistinguishable (data not shown). This suggests that we are in a data rich regime where all of the reweighted ensembles of A$\beta$40 derived from the unbiased a99SB-*disp* A$\beta$40 MD ensemble are extremely similar.

We note that for the a99SB-*disp* A$\beta$40 unbiased MD ensemble, with the exception of backbone RDCs, the agreement of unrestrained cross-validating data with experiment continues to improve as the effective ensemble size is decreased beneath 3000 structures, below the selected Kish Ratio threshold of $K$=0.10 (SI Figure 3C). We find that unrestrained cross-validating continues to improve until the effective ensemble size of the reweighted ensemble contains fewer than 70 structures, demonstrating that we could further improve agreement with unrestrained cross-validating data if were willing to obtain a much sparser final reweighted ensemble. This suggests that using a Kish ratio threshold of $K \geq 0.10$ to determine regularization parameters for each data type results in minimal overfitting when all experimental data are used as restraints.

#### Assessing the accuracy of IDP conformational ensembles

IDP conformational ensembles calculated using the maximum entropy reweighting procedure proposed in this work have exceptional agreement with extensive experimental NMR and SAXS datasets that describe the local and global structural properties of IDPs compared to the agreement observed in unbiased MD simulations performed with state-of-the-art force fields.[16,17] We compare the accuracy of reweighted conformational ensembles of A$\beta$40, drkN SH3, ACTR, PaaA2 and $\alpha$-synuclein derived from 30$\mu$s unbiased a99SB-*disp*, C22* and C36m MD simulations in Figure 1, SI Figure 12, Figure 2, Figure 3 and SI Figure 21, respectively.

We observe similar agreement between calculated and experimental data in the reweighted a99SB-*disp*, C22*, and C36m ensembles of A$\beta$40 (Figure 1), ACTR (Figure 2), and drkN SH3 (SI Figure 12). The $\overline{\mathrm{RMSE_N}}$ of reweighted ensembles of A$\beta$40, are 0.73, 0.63, 0.61, for a99SB-*disp*, C22* and C36m, respectively (Figure 1E); the $\overline{\mathrm{RMSE_N}}$ of the reweighted ensembles of ACTR are 0.84, 0.84, 0.85, for a99SB-*disp*, C22* and C36m, respectively (Figure 2C); and the $\overline{\mathrm{RMSE_N}}$ of reweighted ensembles of drkN SH3 are 0.80, 0.95, 0.91, for a99SB-*disp*, C22* and C36m, respectively (SI Figure 12C). All of the reweighted ensembles of these three IDPs are in exceptionally good agreement with experimental NMR and SAXS data, and are in substantially better agreement with experimental data than the most accurate unbiased MD ensembles of these proteins reported in previous benchmark studies.[16,17]

We observe larger differences between calculated and experimental data in reweighted ensembles of PaaA2 (Figure 3) and $\alpha$-synuclein (SI Figure 21). The $\overline{\mathrm{RMSE_N}}$ of reweighted ensembles of PaaA2 are 0.80, 1.39, 1.14, for a99SB-*disp*, C22* and C36m, respectively (Figure 3C), while the $\overline{\mathrm{RMSE_N}}$ of reweighted ensembles of $\alpha$-synuclein are 0.79, 2.46, 1.97, for a99SB-*disp*, C22* and C36m, respectively (SI Figure 21C). Reweighted a99SB-*disp*, C36m and C22* ensembles of PaaA2 have a similar agreement with experimental NMR chemical shift and RDC data (Figure 3C). The agreement between calculated and experimental data of reweighted C22* and C36m $\alpha$-synuclein ensembles is, however, substantially worse than the agreement observed in the reweighted a99SB-*disp* ensemble (SI Figure 21C).

The agreement with experimental SAXS data is particularly poor in reweighted C22* and C36m ensembles of PaaA2 and $\alpha$-synuclein. The C22* and C36m MD ensembles of these IDPs are substantially more compact than the a99SB-*disp* ensembles both before and after reweighting (Figure 3, SI Figure 18, SI Figure 21, SI Figure 23). Reweighting C22* and C36m ensembles of PaaA2 and $\alpha$-synuclein only produces a marginal improvement in their agreement with the experimental SAXS data (Figure 3A, SI Figure 21A). This results from the fact that unbiased C36m and C22* ensembles of PaaA2 and $\alpha$-synuclein contain almost no conformations with a $R_g$ as large as the experimental $R_g$ determined from SAXS[16,18,19] (SI Figure 18, SI Figure 23). This places an upper limit on the agreement that can be obtained from reweighting regardless of the specified Kish ratio threshold. Reweighted C36m and C22* ensembles of $\alpha$-synuclein also have substantially worse agreement with experimental NMR chemical shift, RDC, and PRE data than the reweighted a99SB-*disp* ensemble, due to an overestimation of $\beta$-sheet content and overly compact ensemble dimensions (SI Figures 21-24).

We present detailed leave-one-out cross validation analyses of reweighting calculations performed on a99SB-*disp* MD ensembles of A$\beta$40, drkN SH3, ACTR, PaaA2 and $\alpha$-synuclein in SI Figure 3, SI Figure 11, SI Figure 7, SI Figure 16 and SI Figure 20, respectively. We observe similar global improvements in the agreement between calculated and experimental data in all proteins, illustrating the robustness of the reweighting protocol proposed here on a range of IDPs. We note one exception in the case of the protein ACTR, where the agreement between calculated and experimental SAXS data obtained using all experimental data as restraints is slightly (6%) worse than the agreement observed in the unbiased a99SB-*disp* MD ensemble of ACTR (SI Figure 7). We also observe that the RMSE between calculated and experimental SAXS data is 18% worse than the RMSE of the unbiased a99SB-*disp* MD ensemble when SAXS data is excluded as cross-validating data. This demonstrates some incompatibility in the agreement between experimental NMR data (backbone chemical shifts, backbone RDCs and PREs) and SAXS data in the conformations sampled in the unbiased a99SB-*disp* MD ensemble of ACTR.

Remarkably, only three of the fifty experimental data types withheld in leave-one-out cross-validation reweighting calculations performed on a99SB-*disp* MD ensembles of A$\beta$40, drkN SH3, ACTR, PaaA2 and $\alpha$-synuclein were found to have worse agreement with experiment in reweighted ensembles than in unbiased MD ensembles: PRE data measured with a spin-label on residue 59 of drkN SH3 (drkN SH3 PRE-59), ACTR SAXS data and $\alpha$-synuclein backbone $^3J_{HNH\alpha}$ coupling constants (SI Figure 11, SI Figure 7, SI Fig 20).

Similarly, only five data types withheld in leave-one-out cross-validation reweighting tests performed on C36m ensembles had worse agreement with experiment in reweighted ensembles than in unbiased MD ensembles (ACTR PRE-3, ACTR PRE-41, ACTR PRE-61, ACTR SAXS data and $\alpha$-synuclein PRE-103) and only three data types withheld in leave-one-out cross-validation reweighting tests performed on C22* ensembles had worse agreement in reweighted ensembles (drkN SH3 PRE-59, ACTR PRE-41 and $\alpha$-synuclein PRE-103). Together, these results demonstrate that the automated maximum entropy reweighting protocol introduced in this investigation to refine IDP ensembles improves agreement with extensive experimental datasets from NMR spectroscopy and SAXS with minimal overfitting.

#### Comparing IDP conformational ensembles derived from different force fields

The populations of $\alpha$-helices in unbiased and reweighted ensembles of ACTR and PaaA2 are shown in Figure 2 and Figure 3, respectively. The populations of $\alpha$-helices in unbiased and reweighted ensembles of A$\beta$40, drkN SH3 and $\alpha$-synuclein are shown in SI Figure 4, SI Figure 12, and SI Figure 21, respectively. We observe highly consistent populations of moderately stable helices (populations of 20%-40%) in the reweighted ensembles of ACTR and drkN (Figure 2, SI Figure 12). In the case of PaaA2, we observe large differences in the locations and populations of helical elements in unbiased MD ensembles that become substantially more similar after reweighting (Figure 3). We note that there are strong signals for the helical elements of drkN SH3, ACTR and PaaA2 in experimental NMR chemical shift, RDC, and NOE data and that the populations of helical conformations observed in our reweighted ensembles are in excellent agreement with helical populations directly estimated from experimental data.[18,20–22]

The populations of $\beta$-sheets in unbiased and reweighted ensembles for A$\beta$40, drkN SH3, ACTR, PaaA2 and $\alpha$-synuclein are shown in Figure 1, SI Figure 13, SI Figure 8, SI Figure 17 and SI Figure 22 respectively. We observe that all reweighted ensembles of A$\beta$40 converge to very similar secondary structure populations, with residues 15-20 and residues 30-35 forming $\beta$-sheets with populations of 20%-40% (Figure 1). We observe several highly populated $\beta$-sheets (populations of 40%-95%) in unbiased C36m and C22* MD ensembles of drkN SH3 and $\alpha$-synuclein that produce notable discrepancies with experimental NMR data (SI Figure 12, SI Figure 13, SI Figure 21, SI Figure 22). The populations of these $\beta$-sheets are only partially decreased upon reweighting. The unbiased C36m MD ensemble of drkN SH3 contains $\beta$-sheets between by residues 1-10 and residues 20-30 with populations of 40%-80%. These populations are reduced to 20%-40% after reweighting with all experimental data. The reweighted C36m drkN SH3 ensemble, however, has substantially worse agreement with NMR chemical shifts and $^3J_{HNH\alpha}$ scalar couplings than the reweighted a99SB-*disp* drkN SH3 ensemble, which contains very little residual $\beta$-sheet structure. We observe similarly modest reductions in spurious populations of $\beta$-sheets in reweighted C22* and C36m ensembles of $\alpha$-synuclein (SI Figure 21, SI Figure 22).

To determine if the persistence of spurious $\beta$-sheet populations is the result of our selected Kish ratio threshold of $K = 0.10$ we computed reweighted C22* and C36m ensembles of drkN SH3 and $\alpha$-synuclein using a Kish ratio threshold of $K = 0.01$. We observed similar $\beta$-sheet populations in these reweighted ensembles, suggesting that the persistence of these $\beta$-sheets is likely the result of the relatively weak effect of residual $\beta$-sheet structure on experimental secondary NMR chemical shifts[23,24] and the properties of empirical structure based chemical predictors. Previous work has shown that experimental secondary NMR chemical shifts and structure-based backbone NMR chemical shift predictions from algorithms such as SHIFTX+[25] and SPARTA+[15] are inherently less discriminative of conformations in the $\beta$-sheet region of backbone $\phi/\psi$ Ramachandran space than conformations in the $\alpha$-helical region.[23,24]

We find that our proposed reweigthing procedure only marginally reduces the populations of potentially spurious $\beta$-sheets in reweighted ensembles of drkN SH3 and $\alpha$-synuclein relative to those calculated by X-EISD[26] and ASTEROIDS.[19] We note that these approaches, and other ensemble calculation approaches that generate conformations by statistically sampling backbone dihedral angles, may not sample any hydrogen-bonded $\beta$-sheet conformations in initial conformational pools used for ensemble selection, but rather aim to identify elevated sampling of $\beta$-sheet Ramachandran space relative to statistical coil models. Therefore, populations of $\beta$-sheets observed in reweighted MD ensembles may not be directly comparable to $\beta$-propensities identified in these ensemble calculation approaches. Nevertheless, we caution that the presence of substantially populated $\beta$-sheets in reweighted MD ensembles produced by our reweighting procedure does not necessarily suggest strong experimental evidence for residual $\beta$-sheet populations. We encourage a direct inspection of agreement with experimental data for these $\beta$-sheet elements. We emphasize that an important difference between ensembles obtained from reweighting long timescale MD simulations and ensemble selection approaches such as ASTEROIDS[19,23] or EOM[27] is that the positions of side chains in reweighted MD ensembles are governed by the physics of the underlying force fields, compared to a stochastic sampling of side chain rotamer libraries to place side chains and avoid steric clashes. The positions of sidechains in reweighted MD ensembles may therefore have more physical meaning.

We observe that IDP ensembles derived from MD simulations run with different force fields converge to very similar populations of residual helical structure after reweighting, but IDP ensembles with identical populations of $\alpha$-helices can have large differences in the lengths of helical elements observed in individual conformations. An IDP ensemble containing ten consecutive residues with 50% $\alpha$-helical populations might be constituted of only structures containing a contiguous ten-residue helix or structures with no helical content. Alternatively, this ensemble could contain conformations with short helical elements where the populations of helical turns are relatively uncorrelated from neighboring residues. To obtain a more detailed measure of the similarity of the individual conformations within unbiased and reweighted IDP ensembles, we compared the free energy surfaces of the unbiased and reweighted ensembles of each protein as a function of $R_g$ and the $\alpha$-helical order parameter $S\alpha$. $S\alpha$ is a measure of the number of seven residue fragments in a structure that resemble an ideal helix[28] (See Methods). The free energy surfaces of unbiased and reweighted ensembles of PaaA2 are shown as a function of $R_g$ and $S\alpha$ in SI Figure 18. The free energy surfaces of unbiased and reweighted ensembles of A$\beta$40, drkN SH3, ACTR and $\alpha$-synuclein are shown as a function of $R_g$ and $S\alpha$ in SI Figure 5, SI Figure 14, SI Figure 9 and SI Figure 23, respectively.

We observe dramatic differences in the free energy surfaces of the unbiased a99SB-*disp*, C36m and C22* ensembles of PaaA2 both before and after reweighting (SI Figure 18). The reweighted a99SB-*disp*, C22* and C36m PaaA2 ensembles have similar per-residue populations of helical conformations but have very little overlap when projected using $R_g$ and $S\alpha$ descriptors. For comparison, we plot the $R_g$ and $S\alpha$ values of 50 conformations contained in a protein ensemble database (PED)[29] ensemble of PaaA2 calculated directly from NMR and SAXS restraints (PED00013)[18] as white dots in each free energy surface in SI Figure 18. We observe that the unbiased a99SB-*disp* PaaA2 MD ensemble has two broad free energy minima in the $R_g$ and $S\alpha$ projection. One of these minima overlaps very well with the conformations in PED00013 ensemble, and the depth of this minimum is substantially increased upon reweighting. The overly compact unbiased C22* and C36m MD ensembles of PaaA2 have relatively little overlap with the experimental PaaA2 ensemble in this projection. Upon reweighting, the overlap with more compact conformations in the experimental ensemble increases, but the reweighted C22* and C36m ensembles are substantially more similar to the unbiased MD ensembles from which they were derived than to the experimentally restrained PED ensemble or the reweighted a99SB-*disp* PaaA2 ensemble. Reweighted PaaA2 ensembles clearly do not converge to similar conformational distributions across different force fields.

Free energy surfaces of unbiased and reweighted drkN SH3 ensembles are compared to an experimental ensemble (PED00427) in SI Figure 14. All three unbiased MD ensembles of drkN SH3 have substantial overlap with the experimental ensemble in the $R_g$ and $S\alpha$ projection, and the free energy surfaces of reweighted ensembles are relatively similar. In contrast to PaaA2, reweighted drkN SH3 ensembles appear to converge to similar conformational distributions with similar lengths of helical elements and a similar coupling between helix formation and chain compaction. Free energy surfaces of reweighted and unbiased ensembles of A$\beta$40, ACTR and $\alpha$-synuclein are shown as a function of $R_g$ and $S\alpha$ in SI Figure 5, SI Figure 9 and SI Figure 23, respectively. Reweighted ensembles of A$\beta$40, which have minimal helical content, are fairly similar in this projection. Unbiased ensembles of ACTR have fairly large differences in this projection, and become more similar after reweighting. Unbiased and reweighted $\alpha$-synuclein ensembles have very little overlap in these projections.

We examine the similarity of the populations of intramolecular contacts, or *contact maps*, of unbiased and reweighted ensembles of A$\beta$40, drkN SH3, ACTR, PaaA2 and $\alpha$-synuclein in SI Figure 6, SI Figure 15, SI Figure 10, SI Figure 19 and SI Figure 24, respectively. Unbiased A$\beta$40 ensembles have a similar pattern of intramolecular contacts, defined by contacts between residues 15-20 and residues 30-35, with notable differences in populations. The populations of these contacts, which are driven by transient $\beta$-sheet formation, become more similar after reweighting (SI Figure 6). The populations of intramolecular contacts of drkN SH3 and ACTR are quite similar in unbiased MD ensembles of C36m and C22*, while the more extended unbiased a99SB-*disp* MD ensembles contain fewer intramolecular contacts. Upon reweighting ensembles of drkN SH3 and ACTR with experimental datasets that contain PREs and SAXS data, the contact maps become more similar, and the contact maps of the reweighted ACTR ensembles are particularly similar (SI Figure 10).

The reweighted C36m and C22* ensembles of drkN SH3 contain several intramolecular contacts resulting from $\beta$-sheet structure not observed in the reweighted a99SB-*disp* ensemble (SI Figure 15). The overly compact unbiased C36m and C22* MD ensembles of PaaA2 and $\alpha$-synuclein contain a large number of highly populated intramolecular contacts relative to the more realistically expanded unbiased a99SB-*disp* MD ensembles (SI Figure 19, SI Figure 24). The contacts observed in C36m and C22* PaaA2 and $\alpha$-synuclein ensembles remain highly populated after reweighting, as there are too few extended conformations in the unbiased

MD ensembles to reduce the populations of these contacts.

#### Comparisons to previous work

The reweighting approach presented here builds upon several successful maximum entropy and Bayesian approaches for reweighting conformational ensembles of flexible molecules with experimental data.[1–3,6–8,10] Refining conformational ensembles of flexible molecules with maximum entropy reweighting poses two inherent challenges: i) determining how to balance the strengths of experimental restraints against one's confidence in the initial conformational ensemble, and ii) determining the relative strengths of different types of experimental restraints. Several heuristic and theoretical approaches have been proposed for determining or inferring the strengths of experimental restraints when reweighting ensembles of flexible molecules.[6–10,14,26] When considering experimental datasets with a small number of experimental data types cross-validation or L-curve/elbow methods provide a practical and direct approach. When performing reweighting using a few sets of experimental data it is reasonable and expedient to determine the strength of each set of restraints by identifying the strength where the agreement of calculated cross-validating data with experiment begin to deteriorate.

When reweighting conformational ensembles with data from many different experiments this process becomes less straightforward and one must attempt to normalize the relative RMSE between predicted and experimental values based on estimates of experimental measurement errors and the errors of forward models used to predict experimental data. This is frequently done through the use of $\chi^2$ values obtained by normalizing RMSE calculations with various error models. While it is possible to propose reasonable estimates of experimental measurement and forward model prediction errors for different types of data, these quantities are inherently uncertain and these estimates introduce a degree of subjectivity into reweighting protocols. Additionally, in cases where many experimental data are highly correlated, such as NMR data that report on the distribution of backbone dihedral angles of IDPs (ie. chemical shfits, scalar coupling constants, RDCs), cross-validation may not be a practical approach to determine the relative strengths of restraints.

We have proposed a heuristic approach for determining the relative strengths of experimental restraints that directly builds upon previous work that considers the Kish ratios of reweighted ensembles.[3] This approach is motivated by several practical considerations. One motivation is a desire to objectively compare the accuracy of IDP ensembles derived from unbiased MD simulations run with force fields with different levels of accuracy. If a reweighting approach for calculating IDP ensembles contains adjustable parameters to determine the relative importance of different types of experimental data in a reweighting objective function, it is possible to adjust the relative "scores" or likelihoods of reweighted ensembles derived from different force fields by adjusting these parameters.

A second motivation for using the effective ensemble size of reweighted ensembles to choose the relative strenghts of experimental restraints is a desire to directly compare the structural properties and similarity of reweighted ensembles derived from different force fields. If we choose to use RMSE (or $\chi^2$) thresholds to determine the strengths of experimental restraints, we observe that reweighted ensembles derived from different force fields can achieve similar RMSEs with experimental data with dramatically different effective ensemble sizes. Requiring an overly compact C22* or C36m ensemble to agree with SAXS data within a specified tolerance could require reducing the ensemble to only tens of structures, while a more realistically extended a99SB-*disp* ensemble might contain 1,000s or 10,000s of structures after reweighting. Comparing the structural properties and structural overlap of reweighted ensembles with orders-of-magnitude differences in effective ensemble sizes is less meaningful than comparing ensembles with similar effective ensemble sizes.

An additional motivation for using the effective ensemble size of reweighted ensembles to determine the relative strengths of restraints is our desire to adequately represent the most populated regions of conformational space sampled in unbiased MD simulations in reweighted IDP ensembles. One application of our research efforts to determine accurate atomic resolution ensembles of IDPs is to obtain mechanistic insight into their binding mechanisms with small molecule drugs.[30–35] These dynamic and heterogeneous binding mechanisms are mediated by statistical distributions of backbone conformations and positions of sidechain pharmacophores. As such, IDP ensembles with a small number of conformations that sparsely sample backbone topologies and the relative positions of sidechain atoms are unlikely to be informative for predicting small molecule binding sites or understanding subtle shifts in the conformational ensembles of IDPs that occur upon small molecule binding.[30,31,33–35]

We note a conceptual similarity of the reweighting procedure proposed here with the concept of gentle ensemble refinement in the recently published work of Köfinger and Hummer.[9] In this work, the authors propose an elegant approach to balance the strength of experimental restraints in maximum entropy reweighting against confidence in an initial prior model. They relate the KL divergence between the weights of an initial and reweighted ensemble to an energy uncertainty of the unbiased ensemble, and propose to select the optimal value of restraint strength by relating the expected force field accuracy in the space of experimental observables to the expected energy variance upon reweighting. While motivated by theoretical considerations, this approach is similar in spirit and in practice to the heuristic approach proposed here, where we specify an acceptable limit to changes in the statistical weights of conformations in unbiased ensembles *a priori*.

We draw attention to the small effective ensemble sizes obtained with our proposed reweighting procedure if we utilize agreement with restrained or unrestrained data as the primary criterion to select the strength of the experimental restraints. We observe that the RMSE between calculated and experimental data of both restrained and unrestrained cross-validating data largely improve as the strength of experimental restraints is increased until ensemble sizes become very small (Kish ratio $K < 0.01$), corresponding to effective ensemble sizes of a few hundred structures. We note that several approaches to calculate IDP conformational ensembles with a minimal set of structures, sometimes referred to as *maximum parsimony*[36] approaches, have previously found that ensembles of ∼50-500 structures are required to best reproduce NMR and SAXS data of IDPs.[18,19,21,23,26,37] It appears that the IDP ensembles obtained by reweighting long timscale MD simulations using cross-validation approaches produce ensembles with a similar size to ensembles calculated by minimal ensemble approaches.

We observe that the reweighted ensemble of IDPs obtained in this investigation have similar structural properties to conformational ensembles calculated from NMR and SAXS data using minimal ensemble or maximum parsimony approaches. We find that $\alpha$-helical populations of reweighted drkN SH3 ensembles obtained in this work agree well with the helical populations of drkN SH3 ensembles derived from NMR and SAXS data with approaches such as ENSEMBLE[37] and X-EISD.[26] We also observe that $\alpha$-helical populations in PaaA2 ensembles obtained from our reweighting approach agree reasonably well with populations obtained in a previously reported PaaA2 ensemble[18] derived from NMR and SAXS data using the EOM approach.[27]

### Methods

#### Experimental data used for reweighting

We used previously reported experimental datasets containing different combinations of experimental NMR and SAXS data to reweight simulations of each protein. Backbone NMR chemical shifts[18,20,21,38,39] and residual dipolar couplings (RDCs)[18,21,22,40,41] were used as restraints for all proteins. Three-bond backbone $^3J$ scalar coupling constants were used as restraints for A$\beta$40[42] and $\alpha$-synuclein[43] ensembles. Paramagnetic relaxation enhancements (PREs) were used as restraints for drkN SH3,[37] ACTR,[21] and $\alpha$-synuclein[40,44,45] ensembles. SAXS data were used as restraints for drkN SH3,[46] ACTR,[47] PaaA2,[18] and $\alpha$-synuclein[19] ensembles. We treat the NMR chemical shifts of each backbone nucleus ($C\alpha, C\beta, C, H\alpha, H, N$), each class of $^3J$ scalar coupling constant (ie. $^3J_{HNH\alpha}$ and $^3J_{CC}$) and PREs measured from each nitroxide spin-label as separate experimental *data types*. The experimental datasets used for calculating structural ensembles of A$\beta$40, drkN SH3, ACTR, PaaA2 and $\alpha$-synuclein ensembles contain 7, 10, 11, 8, and 14 experimental data types with 237, 458, 608, 541, and 1293 total experimental restraints, respectively.

#### Calculating experimental observables

All 30$\mu$s MD simulations analyzed here contained 29976 frames, with a spacing of $\sim$1.0ns per frame.[16] Backbone scalar coupling constants were calculated using previously determined Karplus equations.[48,49] RDCs were calculated with PALES[50] using a local alignment window of 15 residues.[22] Backbone chemical shifts were calculated using SPARTA+[15] with MDTraj.[51] The SAXS profiles of each frame were calculated using Pepsi-SAXS[52] following a previously described protocol[53] setting adjustable solvent parameters to $\delta\rho = 3.34\,\text{e/nm}^3$ and $r_0 = 1.68$ $\mathring{A}$. The SAXS profiles were scaled by dividing the intensities by the intensities in the first scattering angle in order to normalize the datasets and set the initial intensity to 1. PREs were calculated using distances between $C\alpha$ atoms as described previously[54]

#### $\alpha$-helical order parameter $S\alpha$

$S\alpha$ describes the similarity of each seven-residue fragment of a protein to an ideal helix.[28] $S\alpha$ is calculated as follows:

$$S_\alpha = \sum_i^N \frac{1 - \left(\frac{\text{RMSD}\alpha_i}{r_0}\right)^8}{1 - \left(\frac{\text{RMSD}\alpha_i}{r_0}\right)^{12}} \tag{12}$$

where RMSD$\alpha_i$ is the root mean square deviation (RMSD) between a seven-residue fragment of a protein (spanning from residue $i$ to residue $i+6$) and an ideal seven residue helix. Eq. 12 functions as switching function which outputs values ranging from 0 (indicating not helical) to 1 (indicating perfectly helical) for each seven-residue segment. The threshold for this function is adjusted by the parameter $\text{r}_0$, which we set to 0.80 Å. Setting the parameter $\text{r}_0 = 0.80$ Å means that a seven-residue segment with RMSD$\alpha_i > 2.5$ Å effectively contributes a value 0 to the $S\alpha$ sum, while a seven-residue segment with RMSD$\alpha_i < 0.5$ Å contributes a value of 1.0 to the $S\alpha$ sum. The value of $S\alpha$ for a protein conformation can therefore be interpreted as a proxy for the number of seven-residue fragments that resemble an ideal helix.

### Supplementary Figures

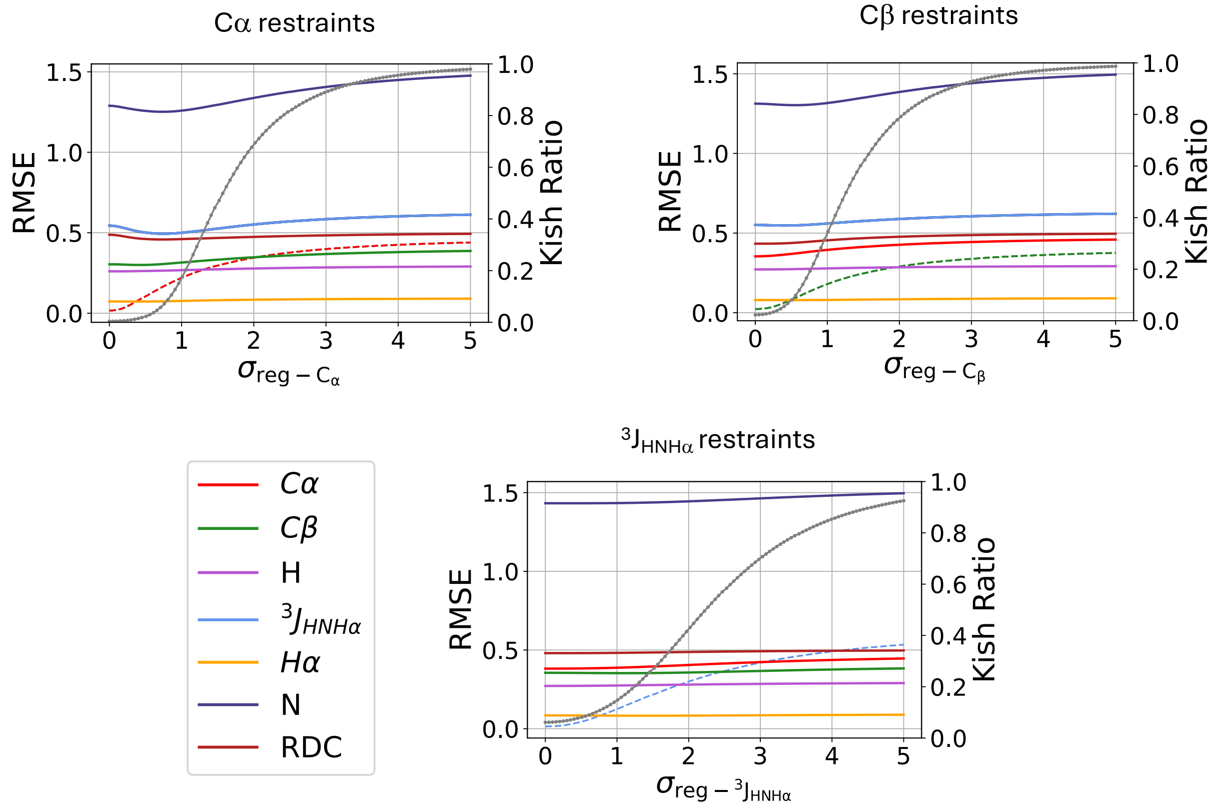


**Supplementary Figure 1: Accuracy of predictions of unrestrained cross-validating data in reweighted A$\beta$40 ensembles obtained by reweighting an a99SB-*disp* MD ensemble with different experimental restraints**. The root mean square error (RMSE) between calculated and experimental data in reweighted A$\beta$40 ensembles are shown as a function of the value of the regularization parameter ($\sigma_{reg}$) of experimental restraints used for reweighting. Experimental data types are indicated by color. The RMSEs of unrestrained cross-validating data are indicated by solid lines, and the RMSE of restrained experimental data is indicated by dotted lines. The Kish ratio of each reweighted ensemble (right y-axis) is indicated by gray dots.

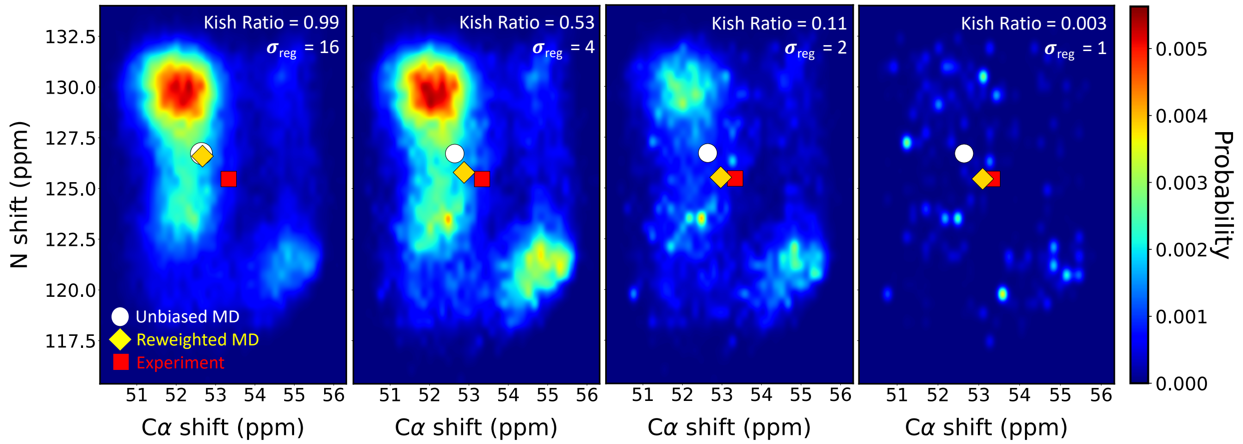


**Supplementary Figure 2: Reweighted IDP ensembles become less representative of the most populated conformational states sampled in unbiased MD simulations as the strength of experimental restraints is increased.** We performed maximum entropy reweighting of a 30$\mu$s unbiased a99SB-*disp* MD ensemble of the ParE2 associated antitoxin protein (PaaA2) using the backbone $C\alpha$ and $N$ NMR chemical shifts of residue ALA38 as restraints. We compare the distribution of the predicted values of these two chemical shifts in each reweighted ensemble as a function of the value of regularization parameter $\sigma_{reg}$ of these restraints. Smaller values of $\sigma_{reg}$ correspond to greater strengths of experimental restraints. The calculated values of the restrained chemical shifts in each reweighted ensemble are shown as yellow diamonds. The calculated values of the chemical shifts in the unbiased MD ensemble are shown as a white circle and the experimental values of the chemical shifts are shown as a red square.

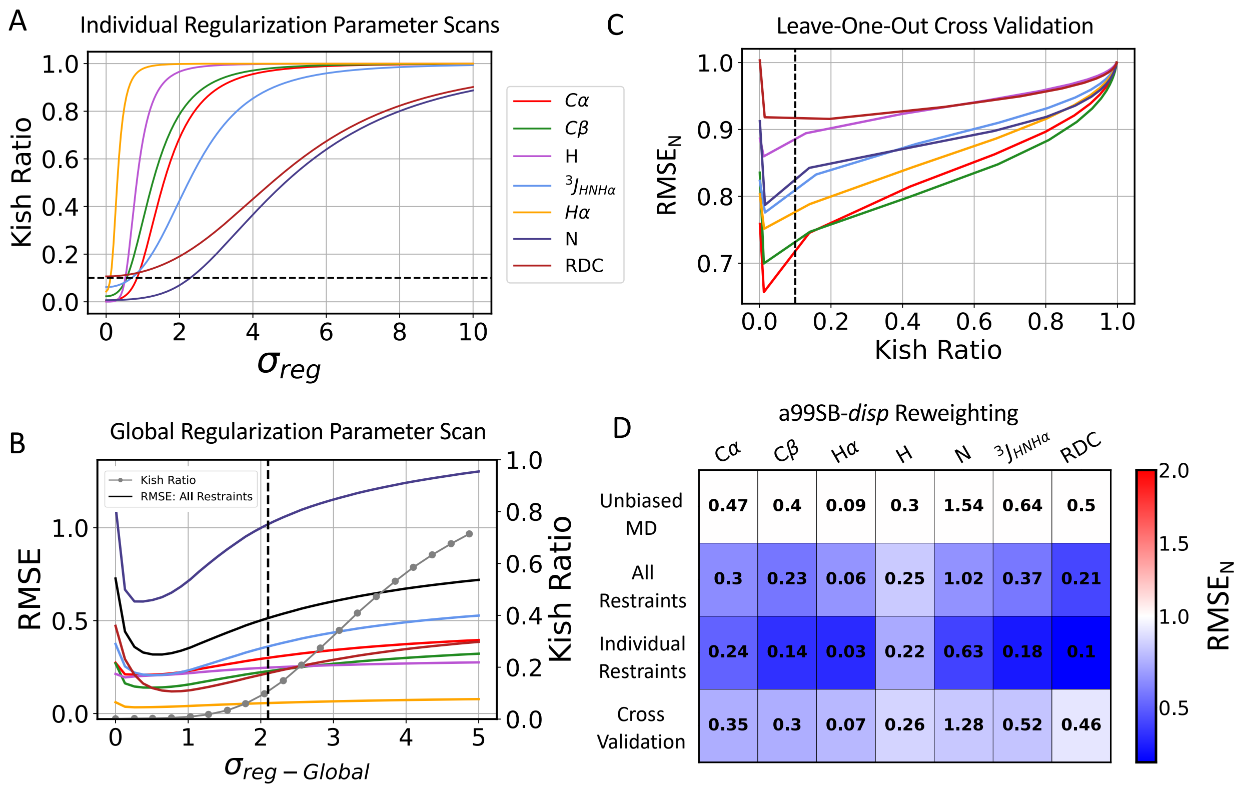


**Supplementary Figure 3: Maximum entropy reweighting of an a99SB-*disp* MD ensemble of A$\beta$40. A)** The effective ensemble size of reweighted ensembles obtained using individual experimental data types as restraints as a function of the value of the regularization parameter $\sigma_{reg}$. **B)** Root mean square error (RMSE) between calculated and experimental data in reweighted ensembles as a function of the value of the global regularization parameter scaling factor ($\sigma_{reg-Global}$). The Kish ratio of each reweighted ensemble is indicated by gray dots. **C)** The normalized RMSE ($\mathrm{RMSE_N}$) of unrestrained data in leave-one-out cross-validation tests where all other experimental data types are used as restraints for reweighting. $\mathrm{RMSE_N}$ values of each data type are normalized by the RMSE observed in the unbiased a99SB-*disp* A$\beta$40 MD ensemble. Black dotted lines indicate the Kish ratio threshold $K = 0.10$ in each plot. **D)** RMSE of experimental data in reweighted ensembles obtained by reweighting the a99SB-*disp* MD ensemble with different combinations of restraints. RMSEs are displayed for the unbiased MD ensemble ("Unbiased MD"), a reweighted ensemble obtained using all experimental data as restraints ("All Restraints"), reweighted ensembles obtained using each individual data type as the only restraint ("Individual Restraints") and reweighted ensembles obtained using all but one data type as restraints ("Cross Validation"). All reweighted ensembles were selected using a Kish ratio threshold of $K = 0.10$. Each square is colored to reflect the value of $\mathrm{RMSE_N}$ relative to the unbiased a99SB-*disp* MD ensemble.

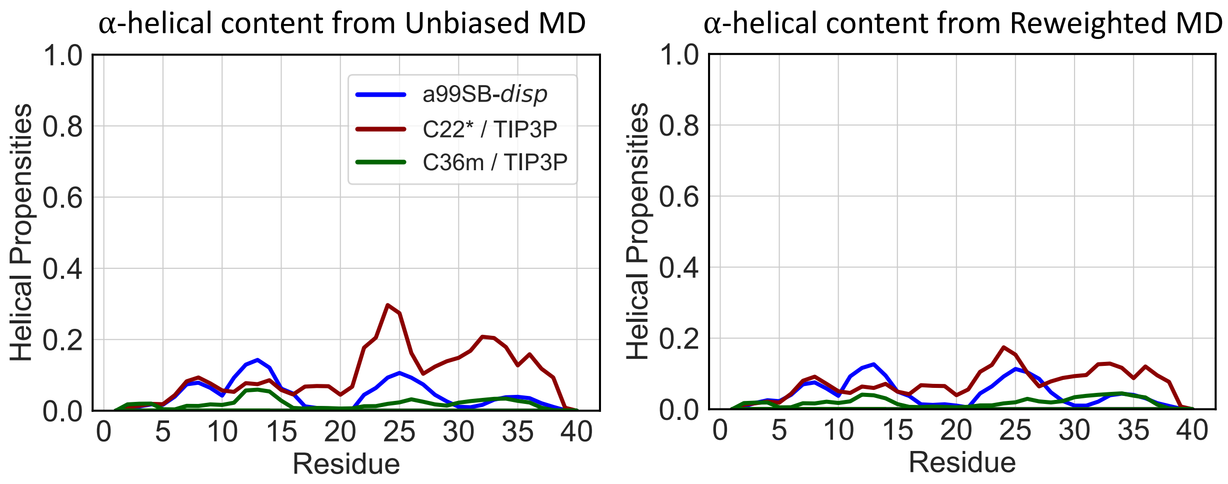


**Supplementary Figure 4: Populations of $\alpha$-helical conformations in unbiased and reweighted MD ensembles of A$\beta$40.**

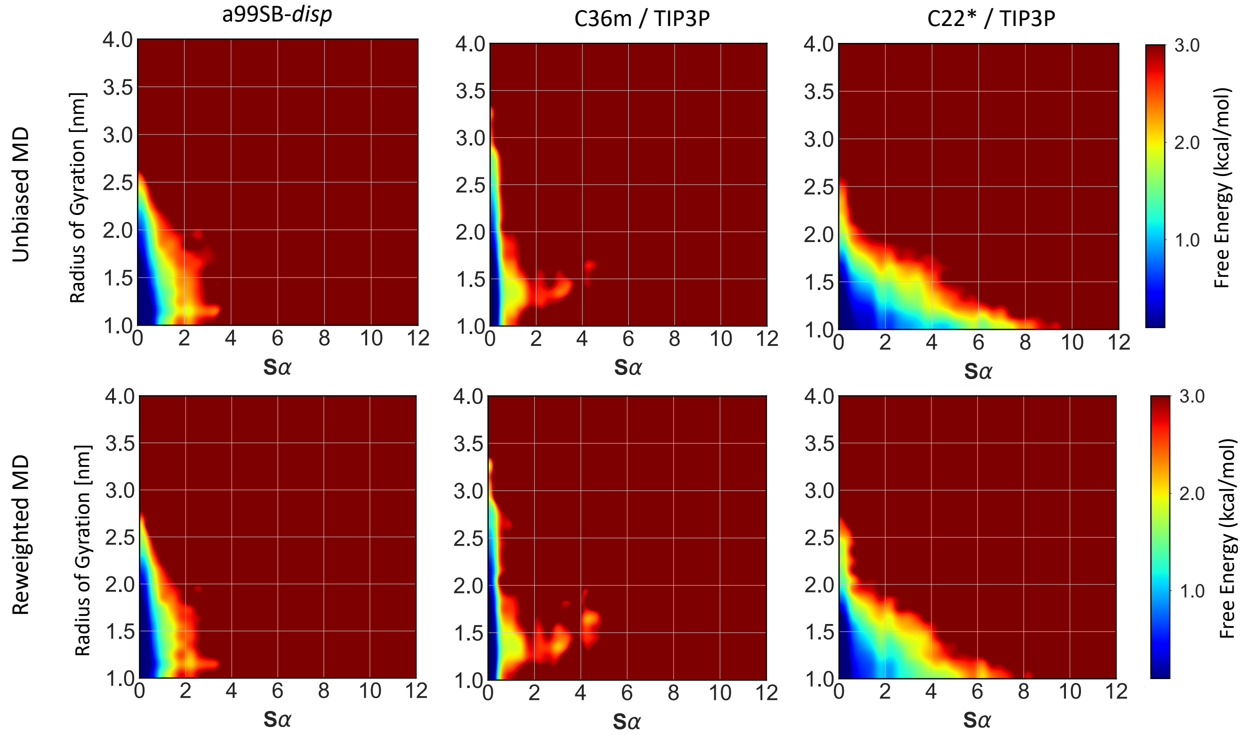


**Supplementary Figure 5: Comparison of free energy surfaces of unbiased and reweighted ensembles of A$\beta$40.** Free energy surfaces of unbiased MD ensembles (top row) and reweighted ensembles (bottom row) of A$\beta$40 are shown as a function of the radius of gyration ($R_g$) and $\alpha$-helical order parameter $S_\alpha$ of each conformation.

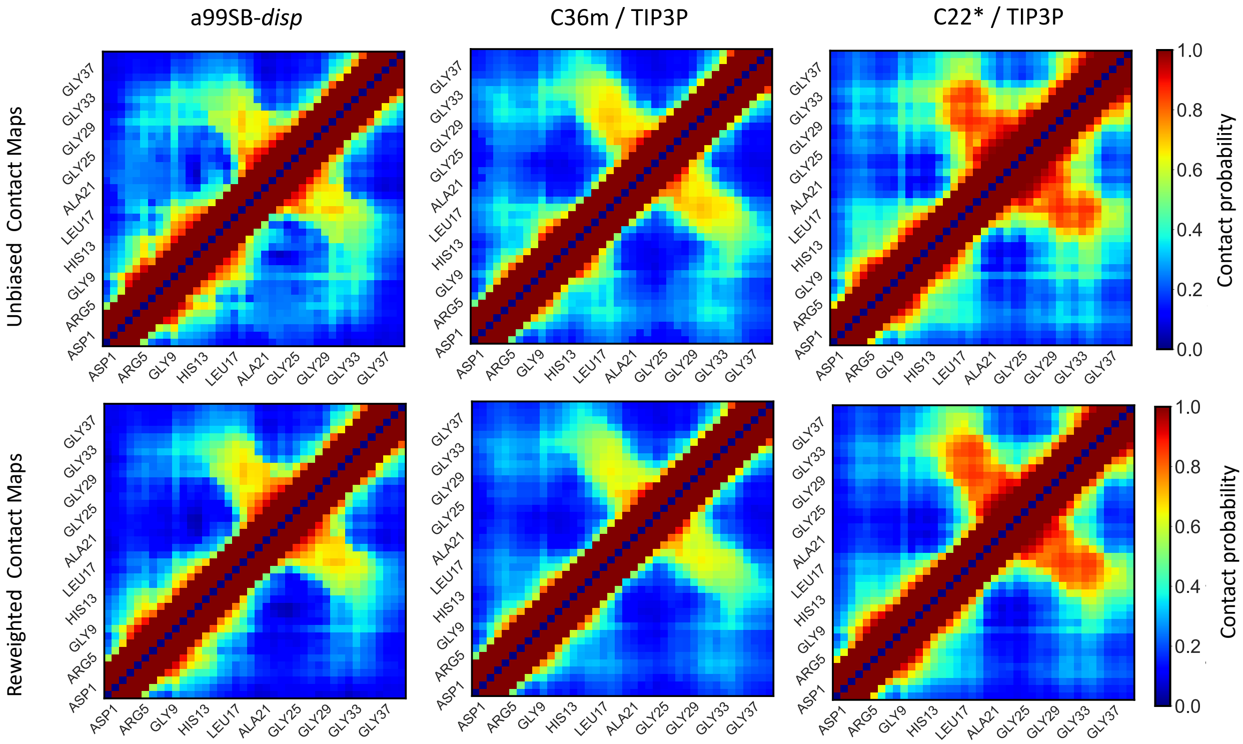


**Supplementary Figure 6: Comparison of the populations of intramolecular contacts in unbiased and reweighted ensembles of A$\beta$40.** Contact maps of unbiased MD ensembles (top row) and reweighted MD ensembles (bottom row) of A$\beta$40. Contacts between residues are defined using a 12 $\AA$ distance threshold between heavy atoms.

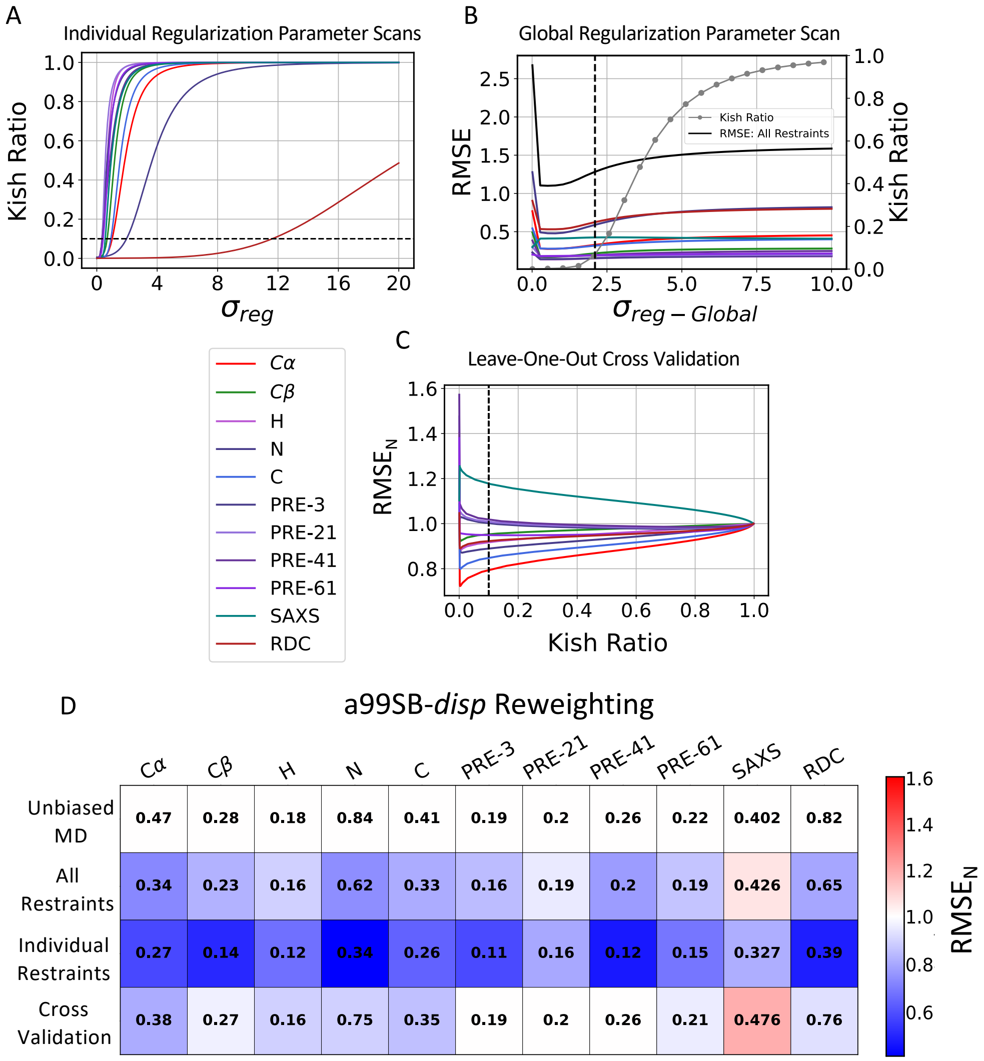


**Supplementary Figure 7: Maximum entropy reweighting of an a99SB-*disp* MD ensemble of ACTR. A)** The effective ensemble size of reweighted ensembles obtained using individual experimental data types as restraints as a function of the value of the regularization parameter $\sigma_{reg}$. **B)** Root mean square error (RMSE) between calculated and experimental data in reweighted ensembles as a function of the value of the global regularization parameter scaling factor ($\sigma_{reg-Global}$). The Kish ratio of each reweighted ensemble is indicated by gray dots. **C)** The normalized RMSE ($\mathrm{RMSE_N}$) of unrestrained data in leave-one-out cross-validation tests where all other experimental data types are used as restraints for reweighting. $\mathrm{RMSE_N}$ values of each data type are normalized by the RMSE observed in the unbiased a99SB-*disp* ACTR MD ensemble. Black dotted lines indicate the Kish ratio threshold $K = 0.10$ in each plot. **D)** RMSE of experimental data in reweighted ensembles obtained by reweighting the a99SB-*disp* MD ensemble with different combinations of restraints. RMSEs are displayed for the unbiased MD ensemble ("Unbiased MD"), a reweighted ensemble obtained using all experimental data as restraints ("All Restraints"), reweighted ensembles obtained using each individual data type as the only restraint ("Individual Restraints") and reweighted ensembles obtained using all but one data type as restraints ("Cross Validation"). All reweighted ensembles were selected using a Kish ratio threshold of $K = 0.10$. Each square is colored to reflect the value of $\mathrm{RMSE_N}$ relative to the unbiased a99SB-*disp* MD ensemble.

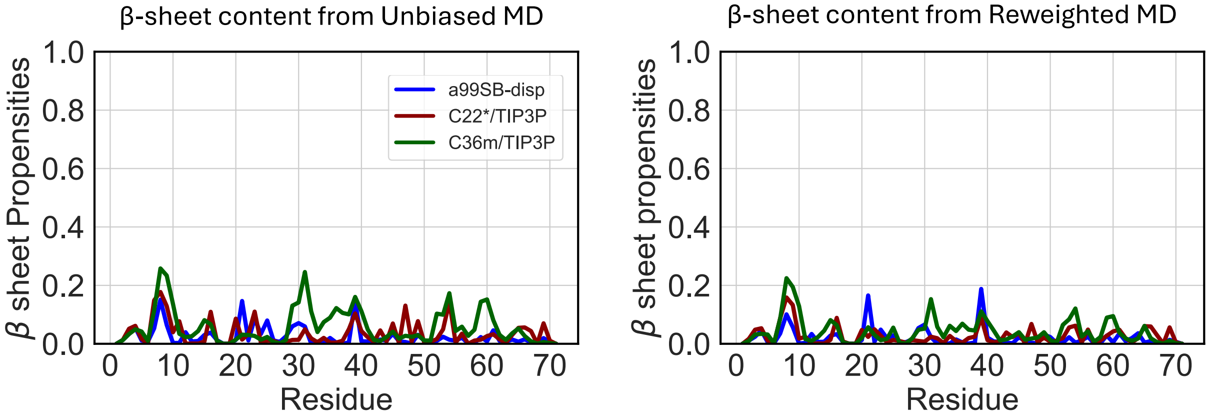


**Supplementary Figure 8: Populations of $\beta$ sheet conformations in unbiased and reweighted MD ensembles of ACTR.**

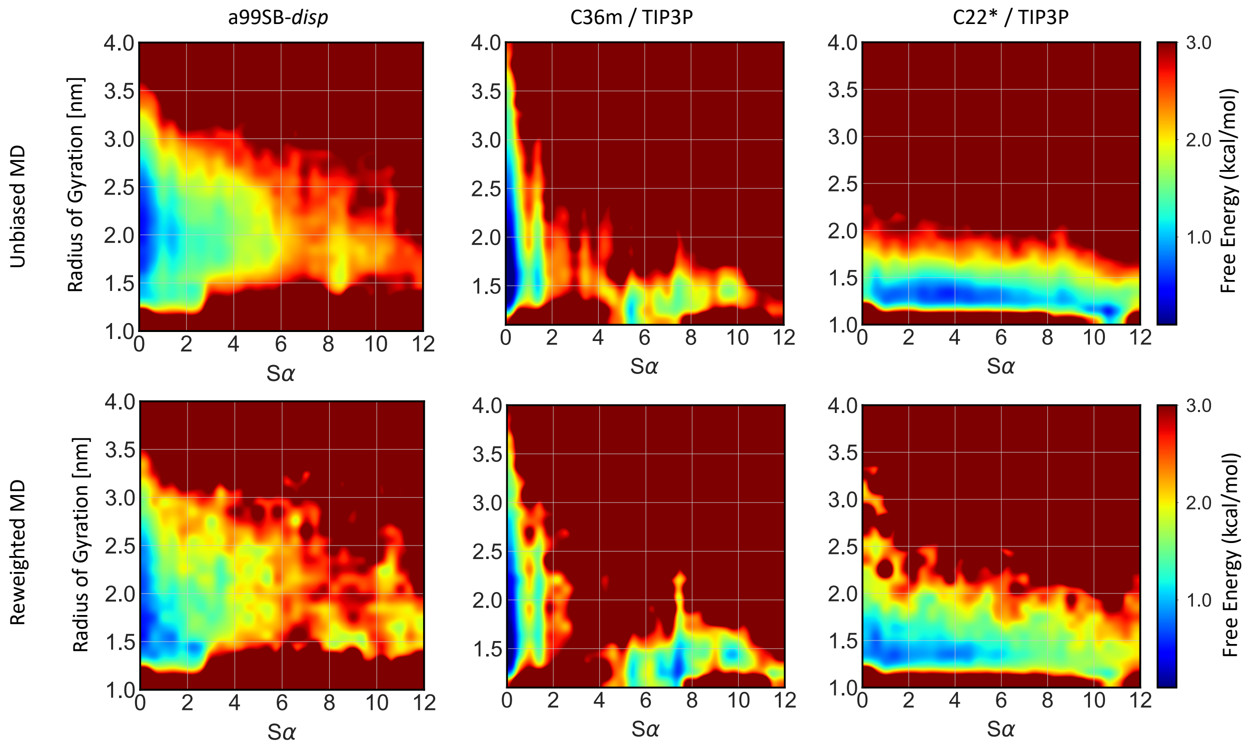


**Supplementary Figure 9: Comparison of free energy surfaces of unbiased and reweighted ensembles of ACTR.** Free energy surfaces of unbiased MD ensembles (top row) and reweighted ensembles (bottom row) of ACTR are shown as a function of the radius of gyration ($R_g$) and $\alpha$-helical order parameter $S_\alpha$ of each conformation.

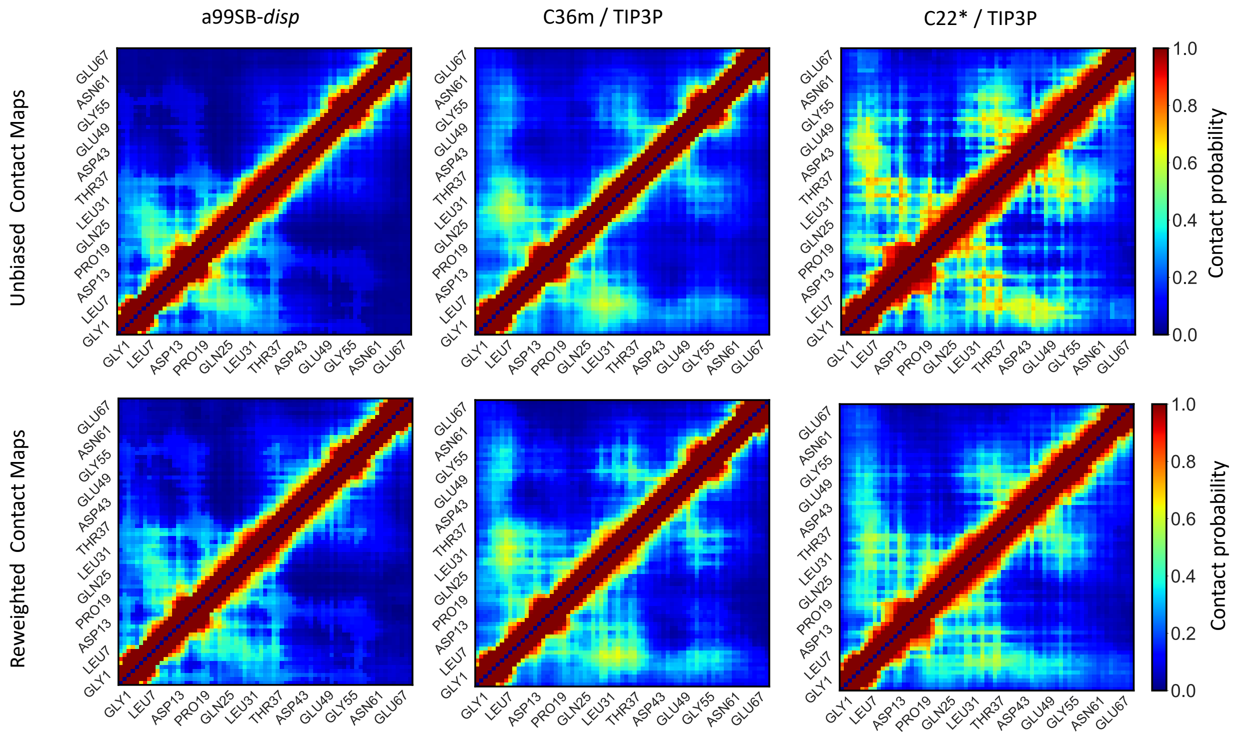


**Supplementary Figure 10: Comparison of the populations of intramolecular contacts in unbiased and reweighted ensembles of ACTR.** Contact maps of unbiased MD ensembles (top row) and reweighted MD ensembles (bottom row) of ACTR. Contacts between residues are defined using a 12 $\mathring{A}$ distance threshold between heavy atoms.

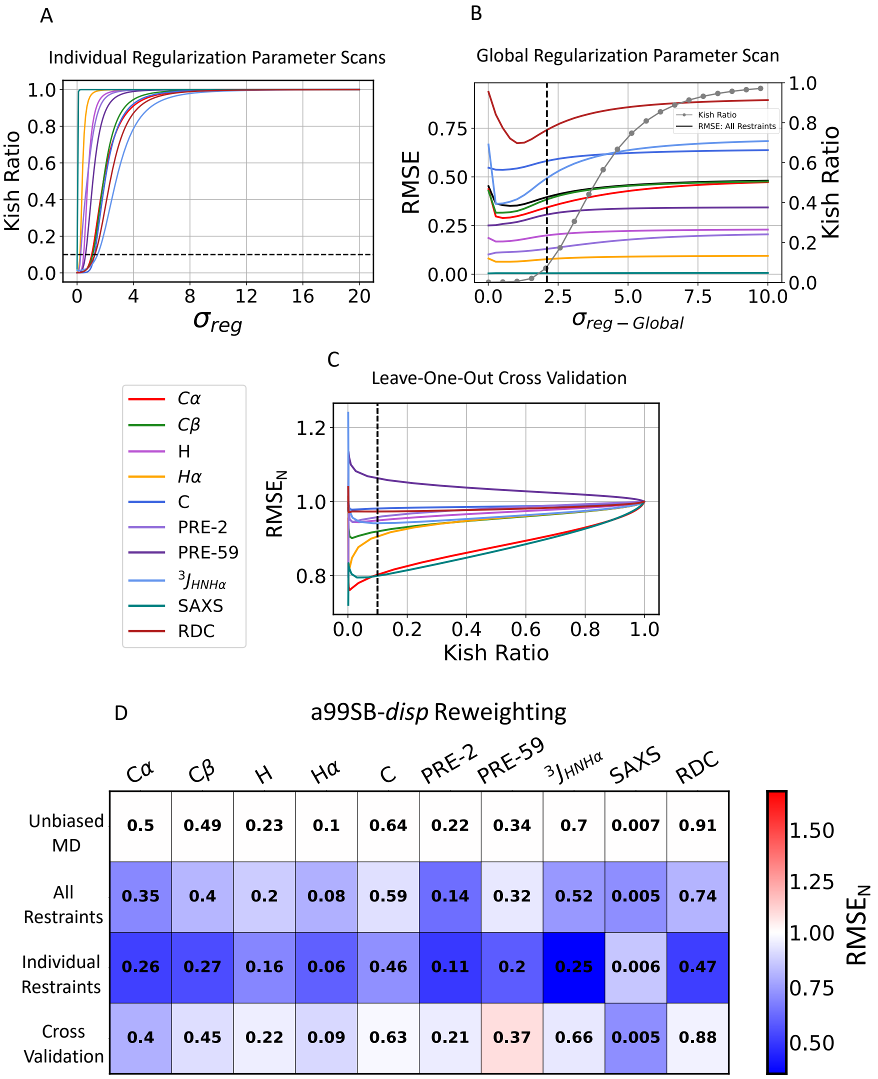


**Supplementary Figure 11: Maximum entropy reweighting of an a99SB-*disp* MD ensemble of drkN SH3. A)** The effective ensemble size of reweighted ensembles obtained using individual experimental data types as restraints as a function of the value of the regularization parameter $\sigma_{reg}$. **B)** Root mean square error (RMSE) between calculated and experimental data in reweighted ensembles as a function of the value of the global regularization parameter scaling factor ($\sigma_{reg-Global}$). The Kish ratio of each reweighted ensemble is indicated by gray dots. **C)** The normalized RMSE ($\mathrm{RMSE_N}$) of unrestrained data in leave-one-out cross-validation tests where all other experimental data types are used as restraints for reweighting. $\mathrm{RMSE_N}$ values of each data type are normalized by the RMSE observed in the unbiased a99SB-*disp* drkN SH3 MD ensemble. Black dotted lines indicate the Kish ratio threshold $K = 0.10$ in each plot. **D)** RMSE of experimental data in reweighted ensembles obtained by reweighting the a99SB-*disp* MD ensemble with different combinations of restraints. RMSEs are displayed for the unbiased MD ensemble ("Unbiased MD"), a reweighted ensemble obtained using all experimental data as restraints ("All Restraints"), reweighted ensembles obtained using each individual data type as the only restraint ("Individual Restraints") and reweighted ensembles obtained using all but one data type as restraints ("Cross Validation"). All reweighted ensembles were selected using a Kish ratio threshold of $K = 0.10$. Each square is colored to reflect the value of $\mathrm{RMSE_N}$ relative to the unbiased a99SB-*disp* MD ensemble.

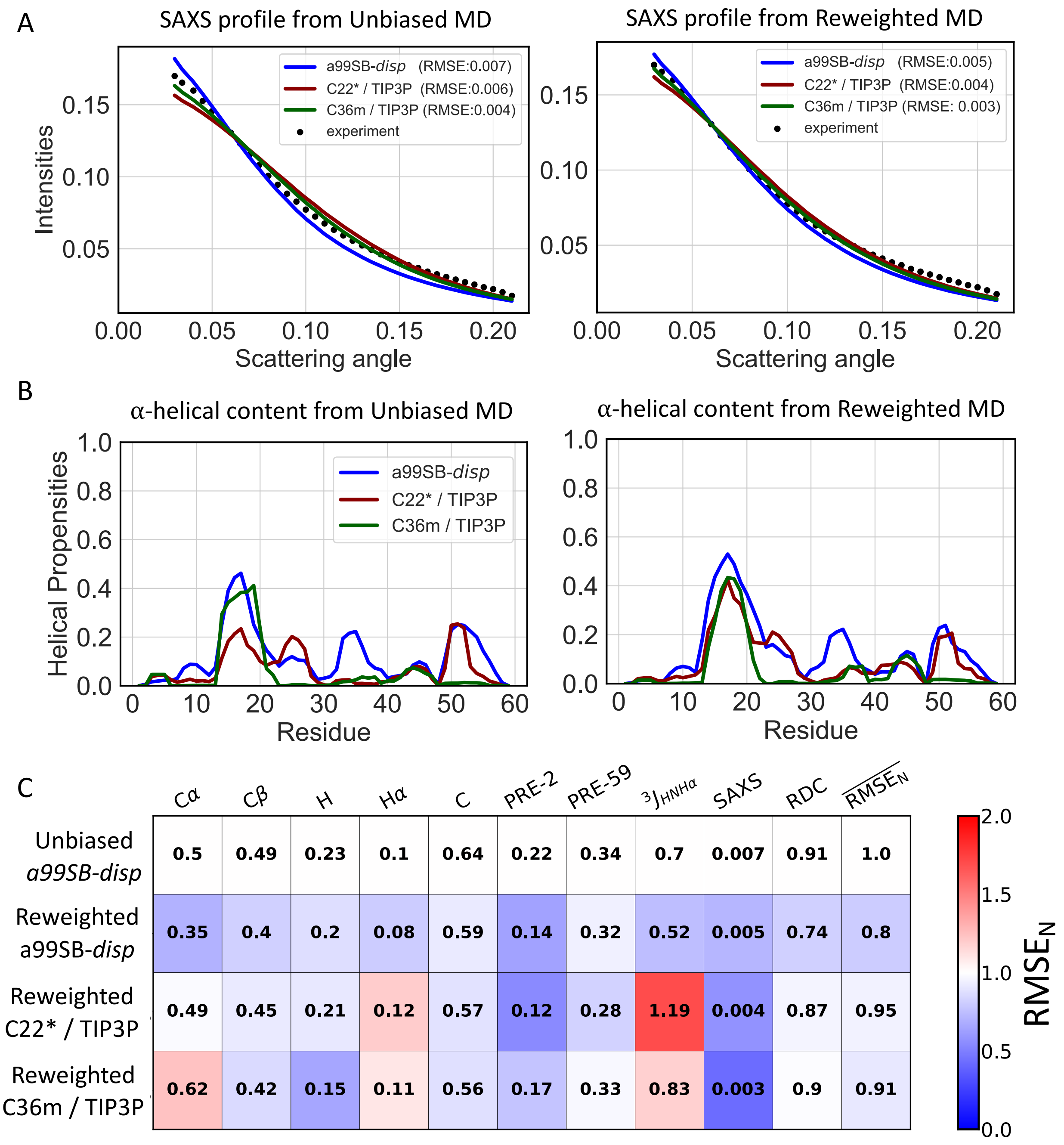


**Supplementary Figure 12: Comparison of unbiased and reweighted MD ensembles of drkN SH3. A)** Comparison of SAXS profiles calculated from unbiased MD ensembles and reweighted MD ensembles of drkN SH3. **B)** Populations of $\alpha$-helical conformations in unbiased MD ensembles and reweighted MD ensembles of drkN SH3. **C)** Comparison of the RMSE between calculated and experimental data in reweighted drkN SH3 ensembles derived from different force fields. Reweighted ensembles were calculated using all experimental data as restraints with a Kish ratio threshold of $K$=0.10. Each square is colored to reflect the value of the normalized RMSE ($RMSE_N$) relative to the unbiased a99SB-*disp* MD ensemble.

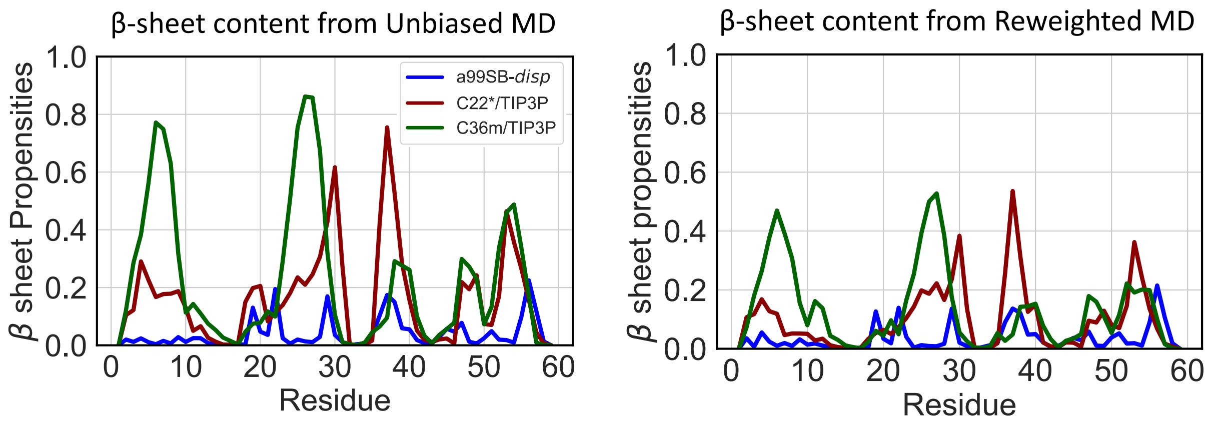


**Supplementary Figure 13: Populations of $\beta$ sheet conformations in unbiased and reweighted MD ensembles of drkN SH3.**

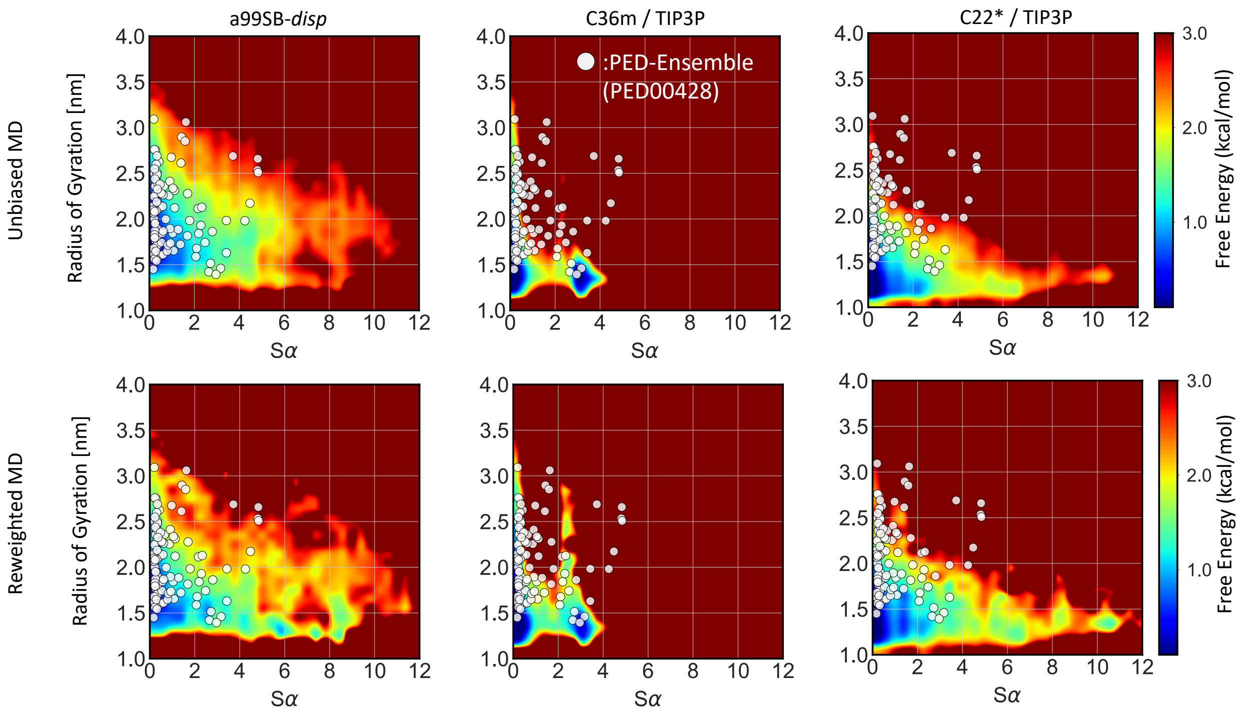


**Supplementary Figure 14: Comparison of free energy surfaces of unbiased and reweighted ensembles of drkN SH3.** Free energy surfaces of unbiased MD ensembles (top row) and reweighted ensembles (bottom row) of drkN SH3 are shown as a function of the radius of gyration ($R_g$) and $\alpha$-helical order parameter $S_\alpha$ of each conformation. The white dots reflect the the $R_g$ and $S_\alpha$ values of structures from a drkN SH3 protein ensemble database (PED) ensemble calculated from experimental NMR data.[55]

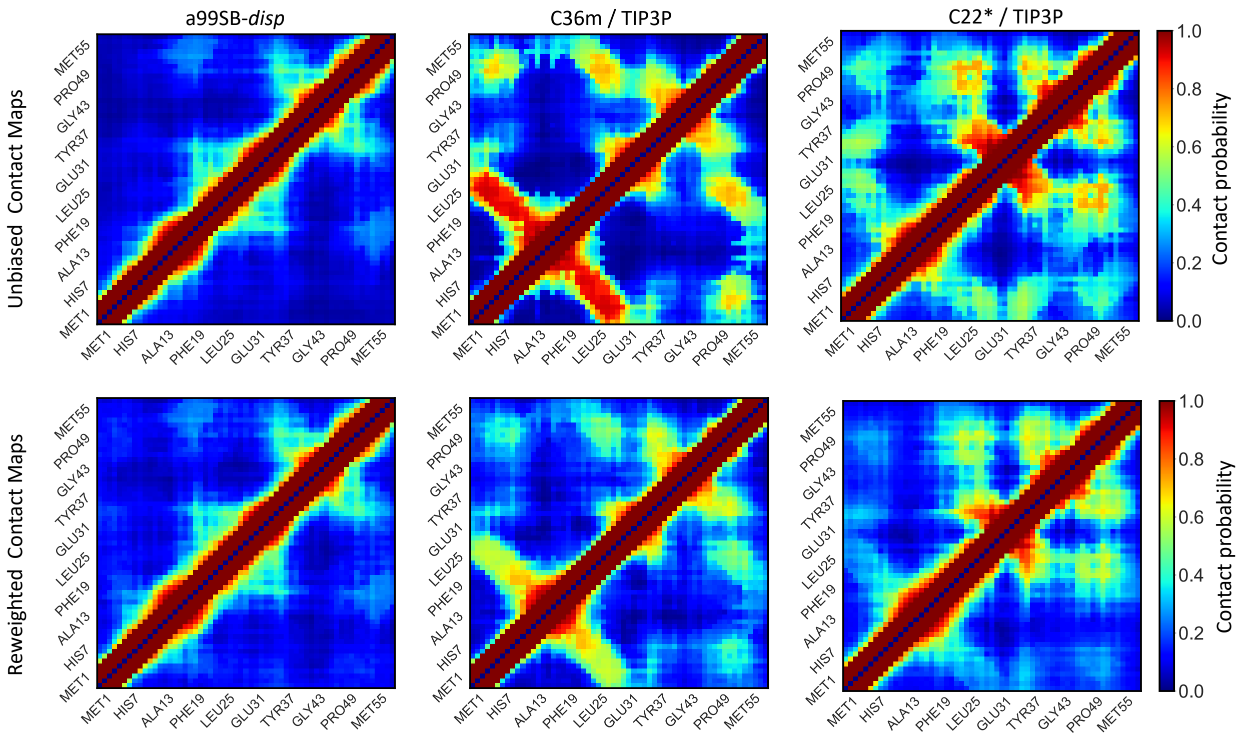


**Supplementary Figure 15: Comparison of the populations of intramolecular contacts in unbiased and reweighted ensembles of drkN SH3.** Contact maps of unbiased MD ensembles (top row) and reweighted MD ensembles (bottom row) of drkN SH3. Contacts between residues are defined using a 12 Å distance threshold between heavy atoms.

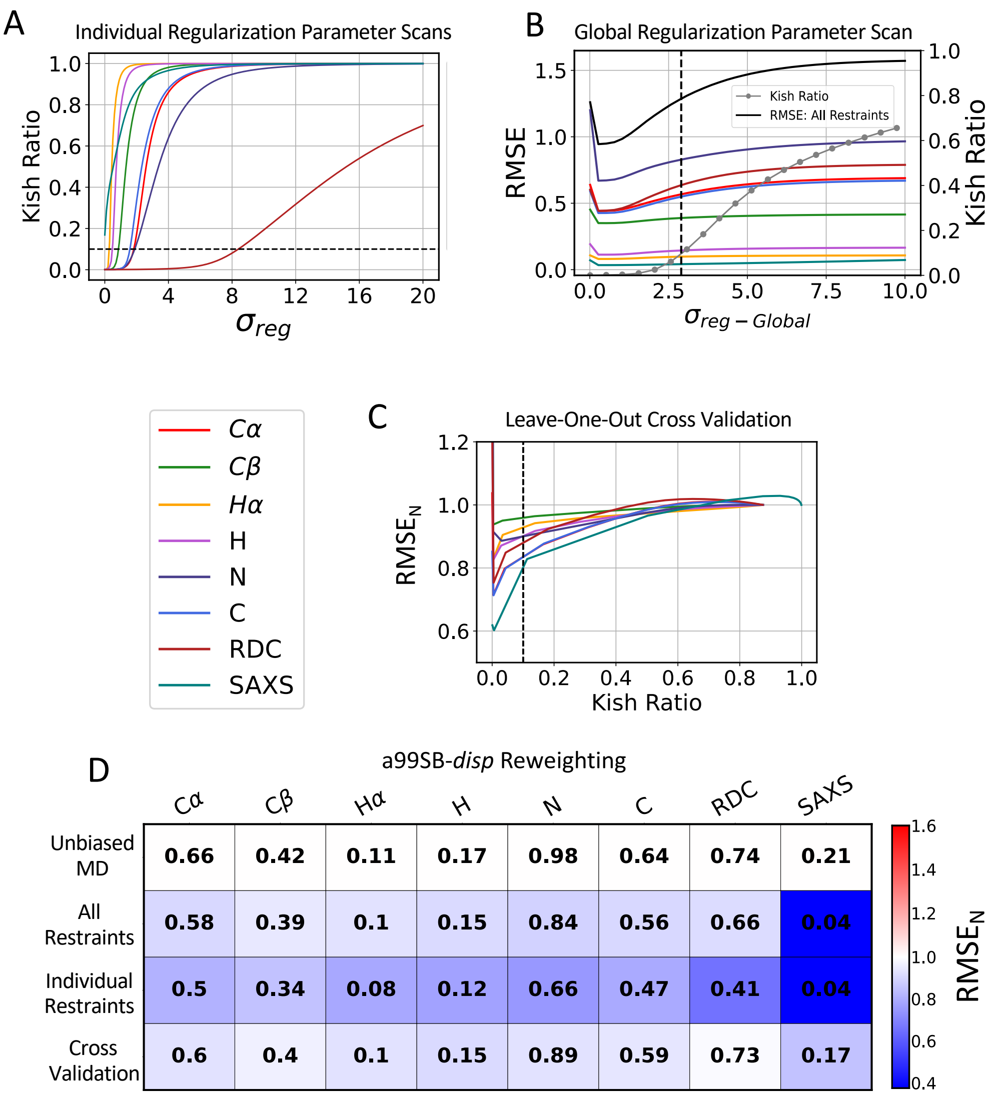


**Supplementary Figure 16: Maximum entropy reweighting of an a99SB-*disp* MD ensemble of PaaA2. A)** The effective ensemble size of reweighted ensembles obtained using individual experimental data types as restraints as a function of the value of the regularization parameter $\sigma_{reg}$. **B)** Root mean square error (RMSE) between calculated and experimental data in reweighted ensembles as a function of the value of the global regularization parameter scaling factor ($\sigma_{reg-Global}$). The Kish ratio of each reweighted ensemble is indicated by gray dots. **C)** The normalized RMSE ($\mathrm{RMSE_N}$) of unrestrained data in leave-one-out cross-validation tests where all other experimental data types are used as restraints for reweighting. $\mathrm{RMSE_N}$ values of each data type are normalized by the RMSE observed in the unbiased a99SB-*disp* PaaA2 MD ensemble. Black dotted lines indicate the Kish ratio threshold $K = 0.10$ in each plot. **D)** RMSE of experimental data in reweighted ensembles obtained by reweighting the a99SB-*disp* MD ensemble with different combinations of restraints. RMSEs are displayed for the unbiased MD ensemble ("Unbiased MD"), a reweighted ensemble obtained using all experimental data as restraints ("All Restraints"), reweighted ensembles obtained using each individual data type as the only restraint ("Individual Restraints") and reweighted ensembles obtained using all but one data type as restraints ("Cross Validation"). All reweighted ensembles were selected using a Kish ratio threshold of $K = 0.10$. Each square is colored to reflect the value of $\mathrm{RMSE_N}$ relative to the unbiased a99SB-*disp* MD ensemble.

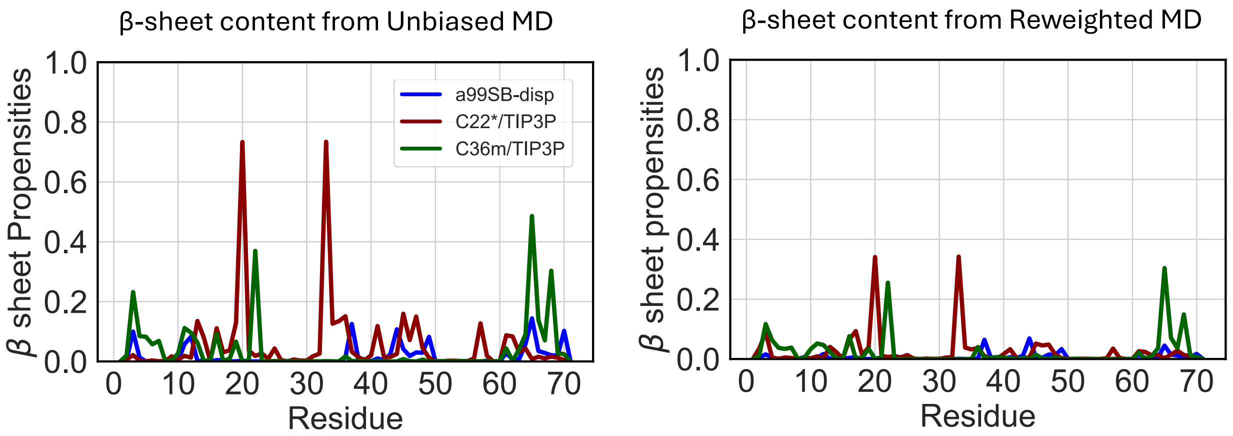


**Supplementary Figure 17: Populations of $\beta$ sheet conformations in unbiased and reweighted MD ensembles of PaaA2.**

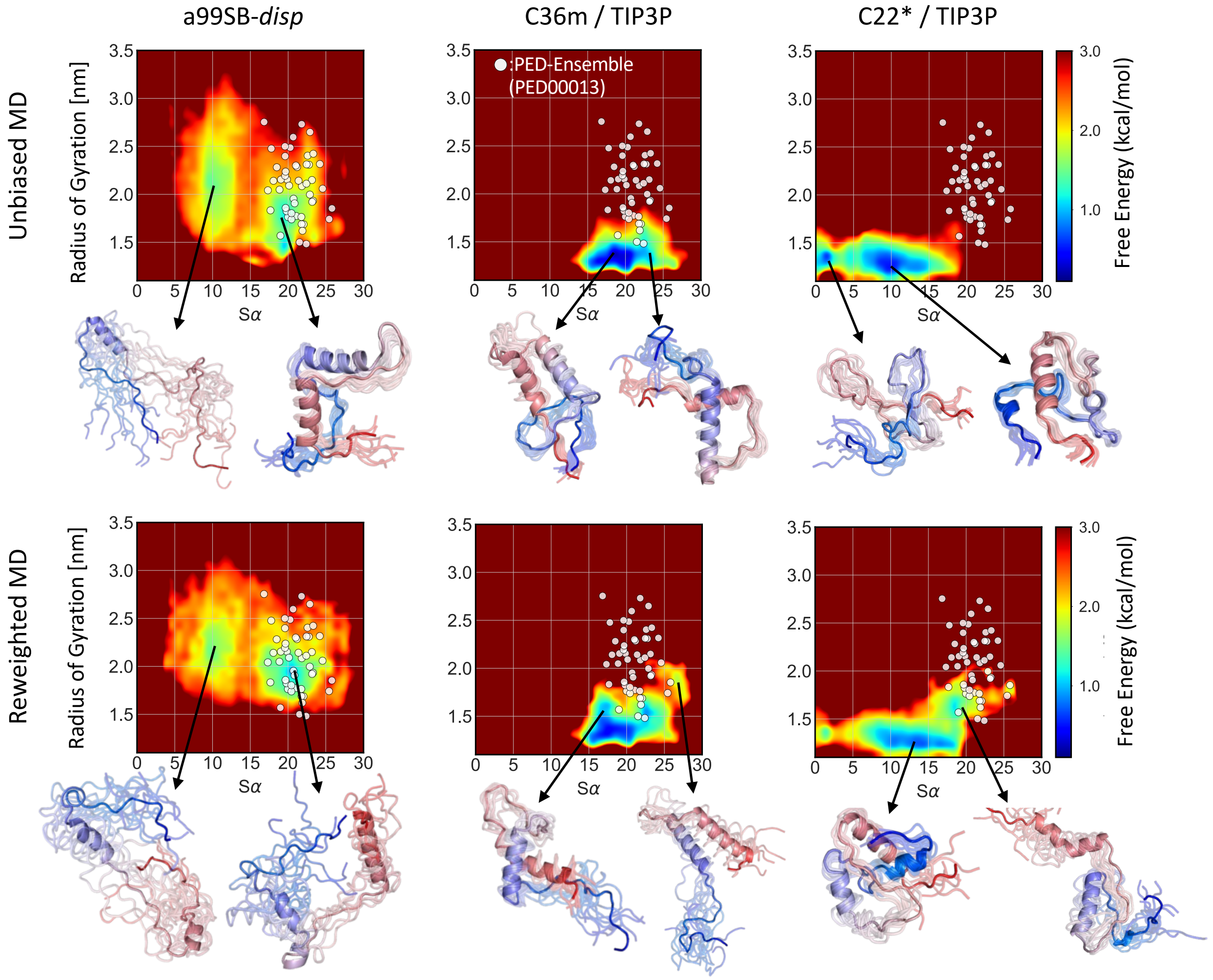


**Supplementary Figure 18: Comparison of free energy surfaces of unbiased and reweighted MD ensembles of PaaA2.** Free energy surfaces of unbiased MD ensembles (top row) and reweighted ensembles (bottom row) of PaaA2 are shown as a function of the radius of gyration ($R_g$) and $\alpha$-helical order parameter $S_\alpha$ of each conformation. The white dots represent the $R_g$ and $S_\alpha$ values of structures from a PaaA2 protein ensemble database (PED) ensemble calculated from experimental NMR and SAXS data.[18] Representative structures are shown for selected free energy basins of each ensemble. Structures are colored with a blue-to-red gradient from the N-terminus to the C-terminus.

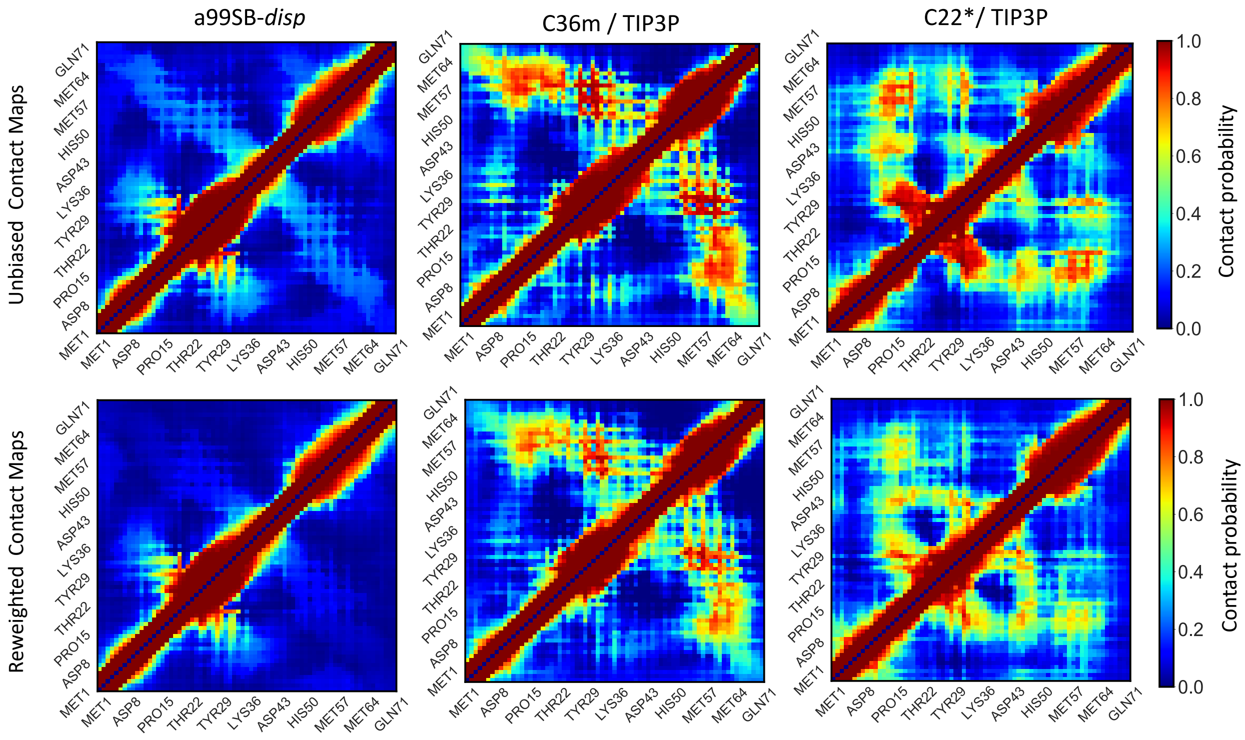


**Supplementary Figure 19: Comparison of the populations of intramolecular contacts in unbiased and reweighted ensembles of PaaA2.** Contact maps of unbiased MD ensembles (top row) and reweighted MD ensembles (bottom row) of PaaA2. Contacts between residues are defined using a 12 $\mathring{A}$ distance threshold between heavy atoms.

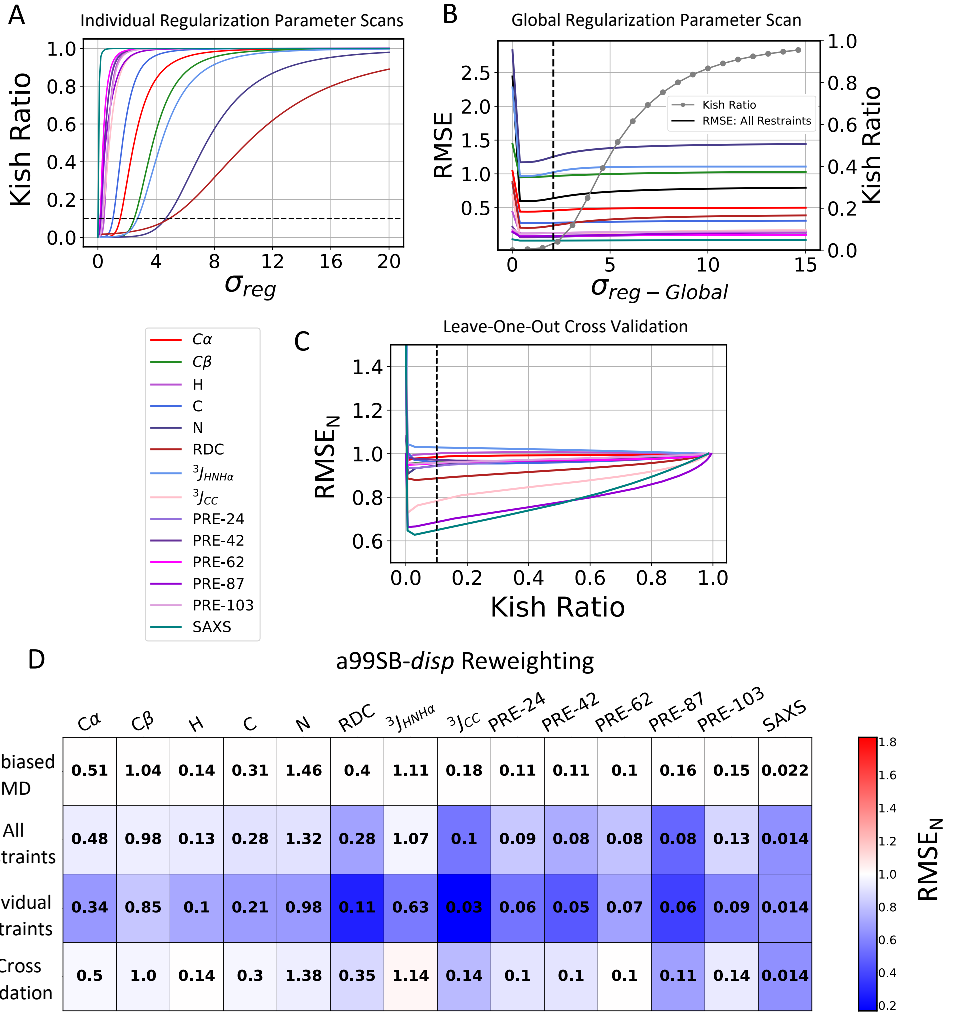


**Supplementary Figure 20: Maximum entropy reweighting of an a99SB-*disp* MD ensemble of $\alpha$-synuclein. A)** The effective ensemble size of reweighted ensembles obtained using individual experimental data types as restraints as a function of the value of the regularization parameter $\sigma_{reg}$. **B)** Root mean square error (RMSE) between calculated and experimental data in reweighted ensembles as a function of the value of the global regularization parameter scaling factor ($\sigma_{reg-Global}$). The Kish ratio of each reweighted ensemble is indicated by gray dots. **C)** The normalized RMSE ($\mathrm{RMSE_N}$) of unrestrained data in leave-one-out cross-validation tests where all other experimental data types are used as restraints for reweighting. $\mathrm{RMSE_N}$ values of each data type are normalized by the RMSE observed in the unbiased a99SB-*disp* $\alpha$-synuclein MD ensemble. Black dotted lines indicate the Kish ratio threshold $K = 0.10$ in each plot. **D)** RMSE of experimental data in reweighted ensembles obtained by reweighting the a99SB-*disp* MD ensemble with different combinations of restraints. RMSEs are displayed for the unbiased MD ensemble ("Unbiased MD"), a reweighted ensemble obtained using all experimental data as restraints ("All Restraints"), reweighted ensembles obtained using each individual data type as the only restraint ("Individual Restraints") and reweighted ensembles obtained using all but one data type as restraints ("Cross Validation"). All reweighted ensembles were selected using a Kish ratio threshold of $K = 0.10$. Each square is colored to reflect the value of $\mathrm{RMSE_N}$ relative to the unbiased a99SB-*disp* MD ensemble.

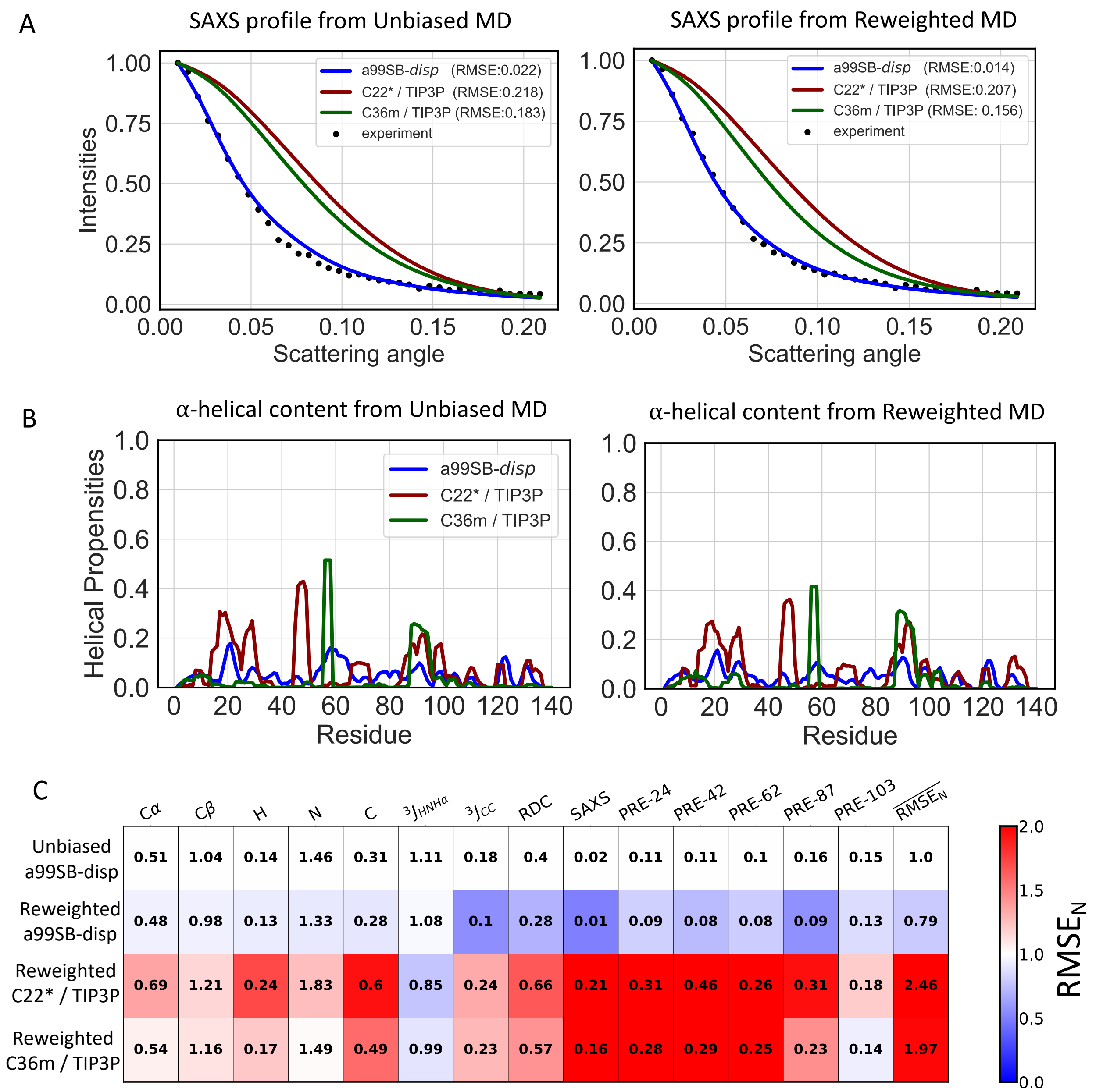


**Supplementary Figure 21: Comparison of unbiased and reweighted MD ensembles of $\alpha$-synuclein. A)** Comparison of SAXS profiles calculated from unbiased MD ensembles and reweighted MD ensembles of $\alpha$-synuclein. **B)** Populations of $\alpha$-helical conformations in unbiased MD ensembles and reweighted MD ensembles of $\alpha$-synuclein. **C)** Comparison of the RMSE between calculated and experimental data in reweighted $\alpha$-synuclein ensembles derived from different force fields. Reweighted ensembles were calculated using all experimental data as restraints with a Kish ratio threshold of $K$=0.10. Each square is colored to reflect the value of the normalized RMSE ($RMSE_N$) relative to the unbiased a99SB-*disp* MD ensemble.

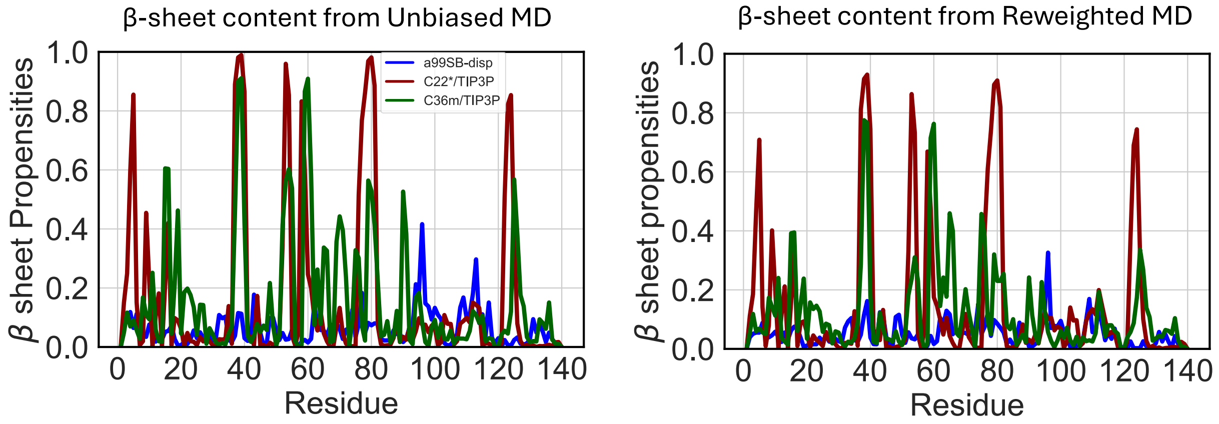


**Supplementary Figure 22: Populations of $\beta$ sheet conformations in unbiased and reweighted MD ensembles of $\alpha$-synuclein**

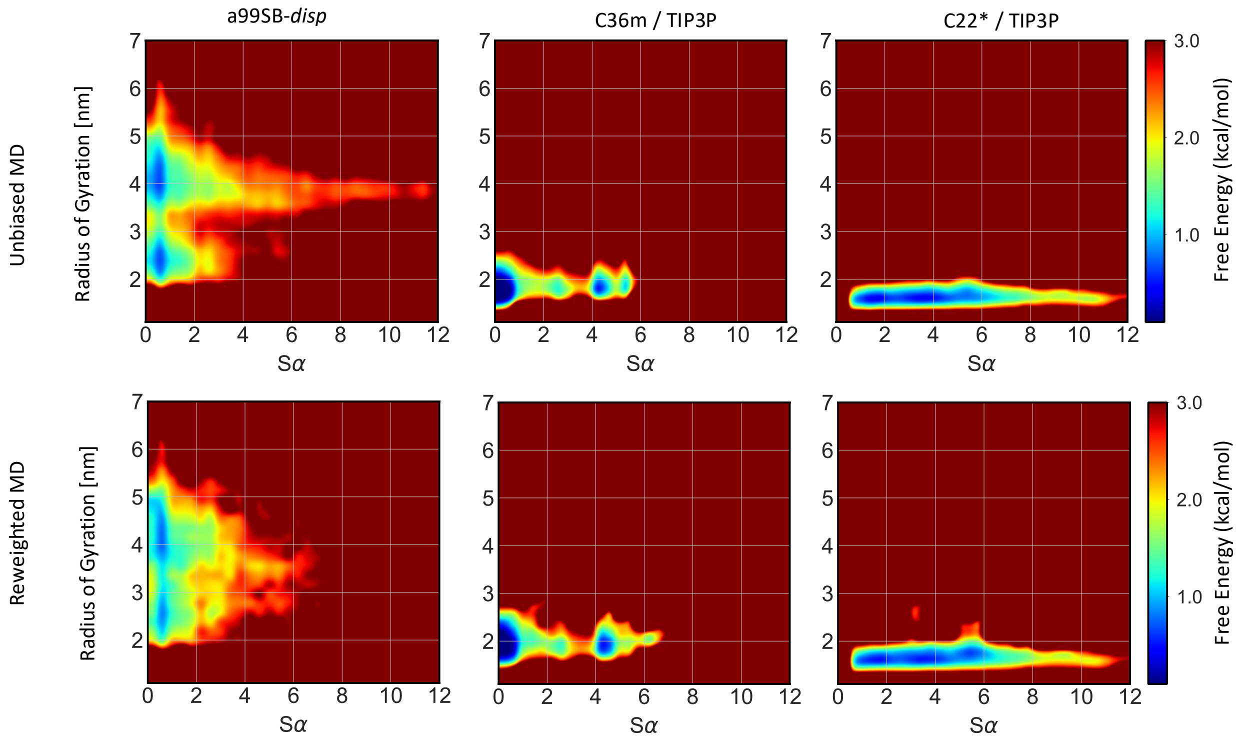


**Supplementary Figure 23: Comparison of free energy surfaces of unbiased and reweighted ensembles of $\alpha$-synuclein.** Free energy surfaces of unbiased MD ensembles (top row) and reweighted ensembles (bottom row) of $\alpha$-synuclein are shown as a function of the radius of gyration ($R_g$) and $\alpha$-helical order parameter $S_\alpha$ of each conformation.

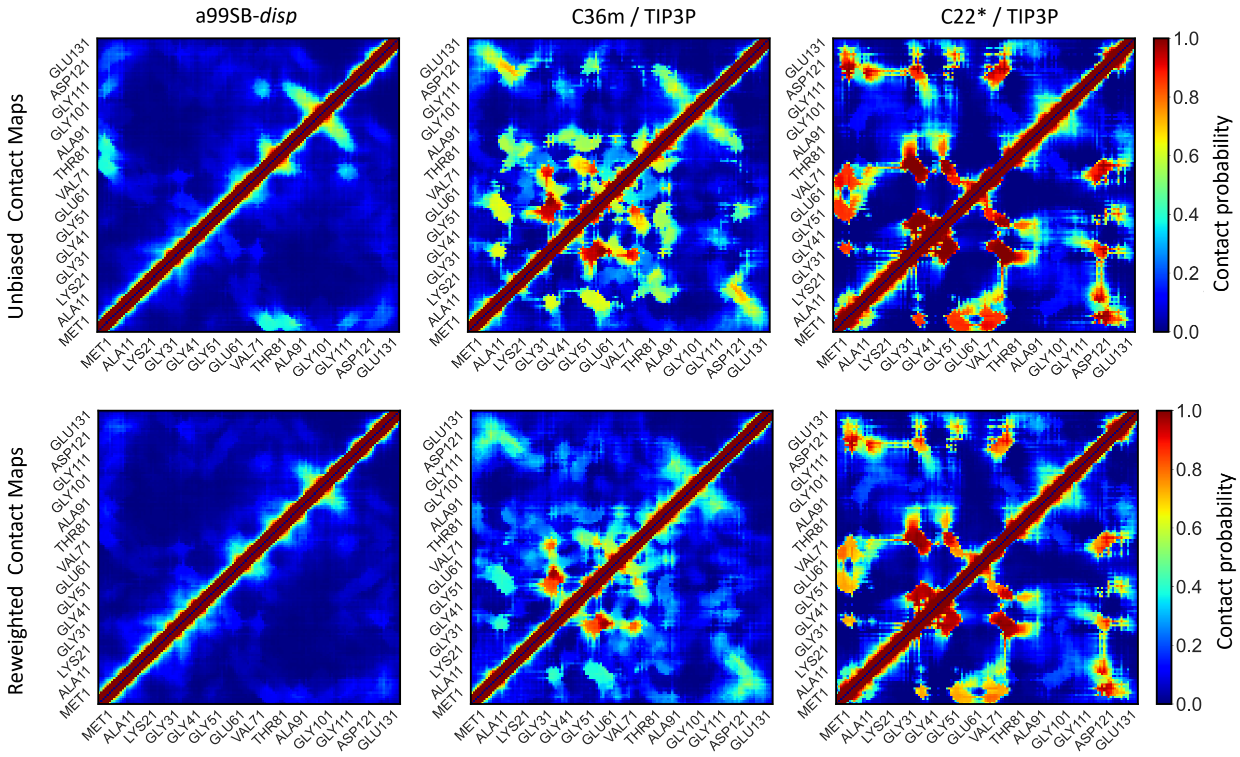


**Supplementary Figure 24: Comparison of the populations of intramolecular contacts in unbiased and reweighted ensembles of $\alpha$-synculein.** Contact maps of unbiased MD ensembles (top row) and reweighted MD ensembles (bottom row) of $\alpha$-synuclein. Contacts between residues are defined using a 12 $\AA$ distance threshold between heavy atoms.

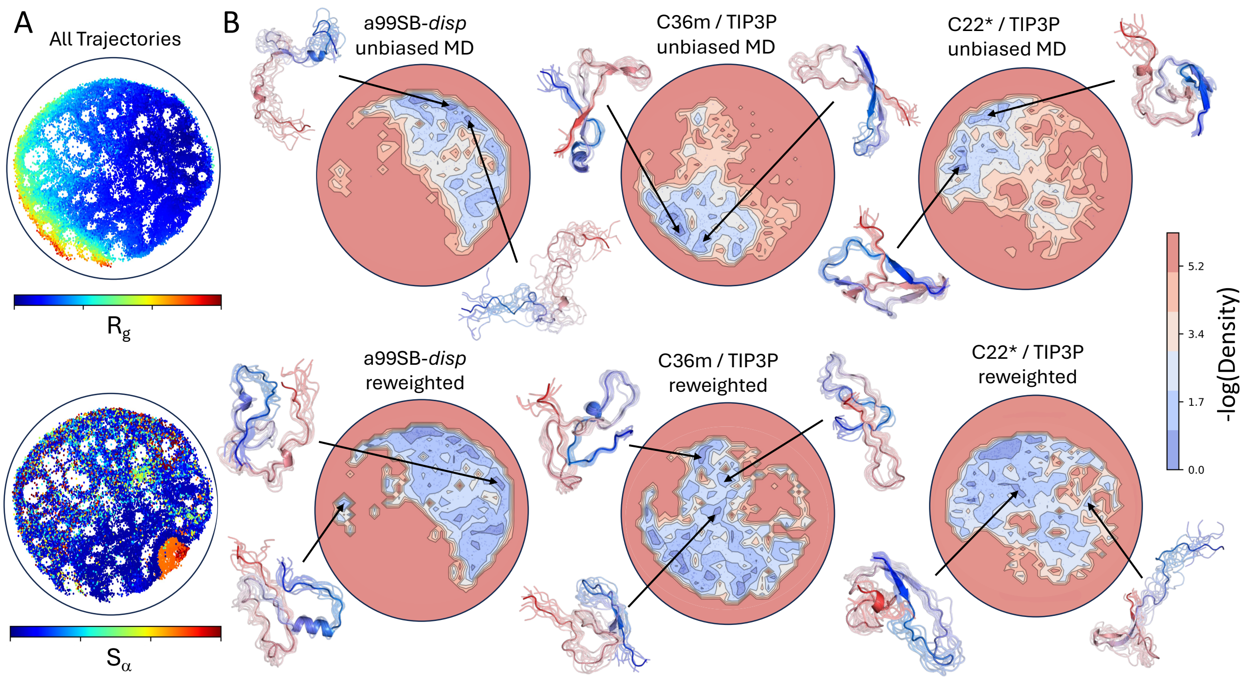


**Supplementary Figure 25: Comparison of energy landscape visualization method (ELViM) projections of unbiased and reweighted MD ensembles of drkN SH3.** A 2D ELViM embedding was calculated using all conformations from unbiased a99SB-*disp*, C22* and C36m MD ensembles of drkN SH3. **A)** The values of the radius of gyration ($R_g$) and the $\alpha$-helical order parameter $S_\alpha$ of each conformation in the unbiased MD ensembles of drkN SH3 projected onto the ELViM latent space. **B)** Comparison of the density of ELViM projections of unbiased and reweighted MD ensembles of drkN SH3. Representative structures are shown for regions of high density in ELViM projections of each ensemble. Structures are colored with a blue-to-red gradient from the N-terminus to the C-terminus.

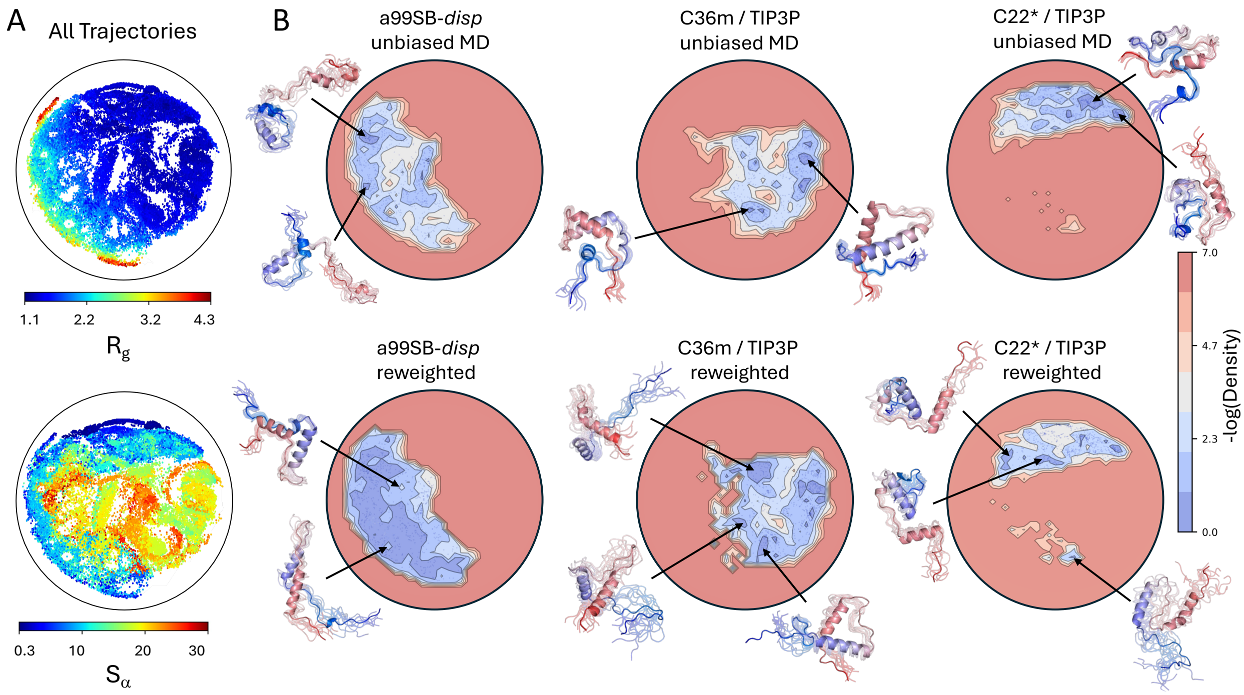


**Supplementary Figure 26: Comparison of energy landscape visualization method (ELViM) projections of unbiased and reweighted MD ensembles of PaaA2.** A 2D ELViM embedding was calculated using all conformations from unbiased a99SB-*disp*, C22* and C36m MD ensembles of PaaA2. **A)** The values of the radius of gyration ($R_g$) and the $\alpha$-helical order parameter $S_\alpha$ of each conformation in the unbiased MD ensembles of PaaA2 projected onto the ELViM latent space. **B)** Comparison of the density of ELViM projections of unbiased and reweighted MD ensembles of PaaA2. Representative structures are shown for regions of high density in ELViM projections of each ensemble. Structures are colored with a blue-to-red gradient from the N-terminus to the C-terminus.

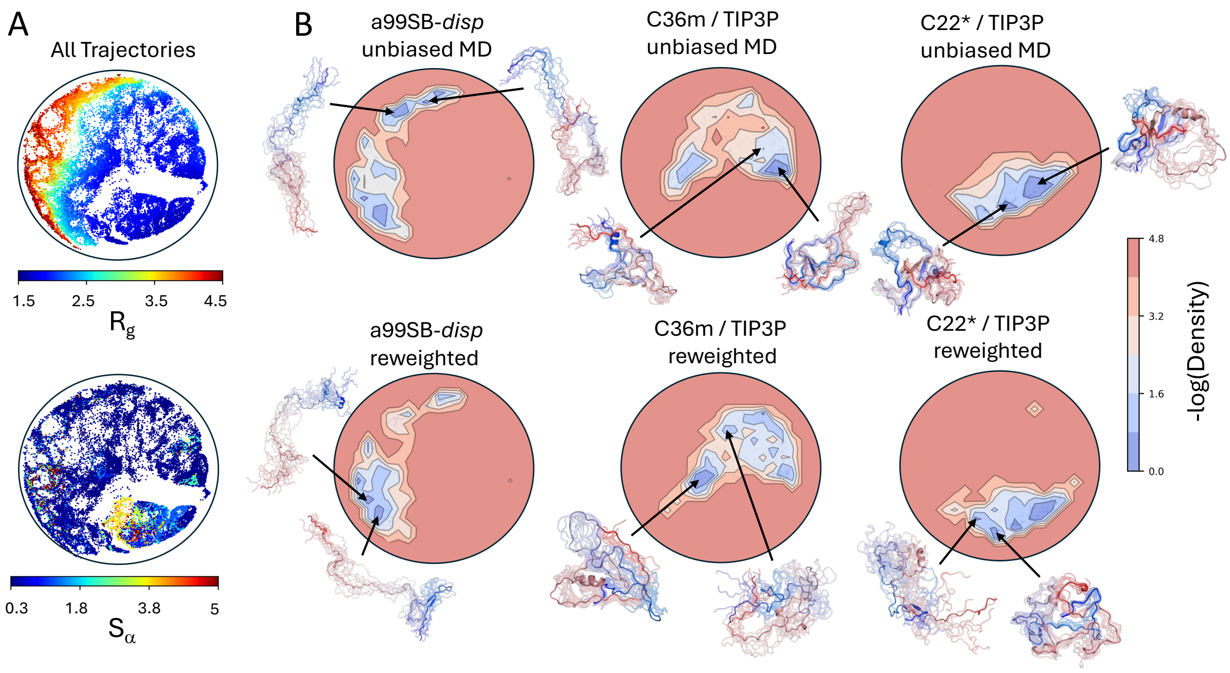


**Supplementary Figure 27: Comparison of energy landscape visualization method (ELViM) projections of unbiased and reweighted MD ensembles of $\alpha$-synuclein.** A 2D ELViM embedding was calculated using all conformations from unbiased a99SB-*disp*, C22* and C36m MD ensembles of $\alpha$-synculein. **A)** The values of the radius of gyration ($R_g$) and the $\alpha$-helical order parameter $S_\alpha$ of each conformation in the unbiased MD ensembles of $\alpha$-synculein projected onto the ELViM latent space. **B)** Comparison of the density of ELViM projections of unbiased and reweighted MD ensembles of $\alpha$-synculein. Representative structures are shown for regions of high density in ELViM projections of each ensemble. Structures are colored with a blue-to-red gradient from the N-terminus to the C-terminus.

(3) Rangan, R.; Bonomi, M.; Heller, G. T.; Cesari, A.; Bussi, G.; Vendruscolo, M. De-

termination of structural ensembles of proteins: restraining vs reweighting. *Journal of chemical theory and computation* **2018**, *14*, 6632–6641.

(13) Kish, L. Survey sampling. 1965.

(51) McGibbon, R. T.; Beauchamp, K. A.; Harrigan, M. P.; Klein, C.; Swails, J. M.; Hernández, C. X.; Schwantes, C. R.; Wang, L.-P.; Lane, T. J.; Pande, V. S. MDTraj:

a modern open library for the analysis of molecular dynamics trajectories. *Biophysical journal* **2015**, *109*, 1528–1532.

(52) Grudinin, S.; Garkavenko, M.; Kazennov, A. Pepsi-SAXS: an adaptive method for rapid and accurate computation of small-angle X-ray scattering profiles. *Acta Crystallographica Section D: Structural Biology* **2017**, *73*, 449–464.

(53) Pesce, F.; Lindorff-Larsen, K. Refining conformational ensembles of flexible proteins against small-angle x-ray scattering data. *Biophysical journal* **2021**, *120*, 5124–5135.

(54) Piana, S.; Donchev, A. G.; Robustelli, P.; Shaw, D. E. Water dispersion interactions strongly influence simulated structural properties of disordered protein states. *The journal of physical chemistry B* **2015**, *119*, 5113–5123.

(55) Teixeira, J. M.; Liu, Z. H.; Namini, A.; Li, J.; Vernon, R. M.; Krzeminski, M.; Shamandy, A. A.; Zhang, O.; Haghighatlari, M.; Yu, L.; others IDPConformerGenerator: a flexible software suite for sampling the conformational space of disordered protein states. *The Journal of Physical Chemistry A* **2022**, *126*, 5985–6003.
